# Multistability and predominant double-positive states in a four node mutually repressive network: a case study of Th1/Th2/Th17/T-reg differentiation

**DOI:** 10.1101/2024.01.30.575880

**Authors:** Atchuta Srinivas Duddu, Elizabeth Andreas, BV Harshavardhan, Kaushal Grover, Vivek Raj Singh, Kishore Hari, Siddharth Jhunjhunwala, Breschine Cummins, Tomas Gedeon, Mohit Kumar Jolly

## Abstract

Elucidating the emergent dynamics of complex regulatory networks enabling cellular differentiation is crucial to understand embryonic development and suggest strategies for synthetic circuit design. A well-studied network motif often driving cellular decisions is a toggle switch - a set of two mutually inhibitory lineage-specific transcription factors A and B. A toggle switch often enables two possible mutually exclusive states - (high A, low B) and (low A, high B) - from a common progenitor cell. However, the dynamics of networks enabling differentiation of more than two cell types from a progenitor cell is not well-studied. Here, we investigate the dynamics of four master regulators A, B, C and D inhibiting each other, thus forming a toggle tetrahedron. Our simulations show that a toggle tetrahedron predominantly allows for co-existence of six ‘double positive’ or hybrid states where two of the nodes are expressed relatively high as compared to the remaining two - (high A, high B, low C, low D), (high A, low B, high C, low D), (high A, low B, low C, high D), (low A, high B, high C, low D), (low A, low B, high C, high D) and (low A, high B, low C, high D). Stochastic simulations showed state-switching among these phenotypes, indicating phenotypic plasticity. Finally, we apply our results to understand the differentiation of naive CD4^+^ T cells into Th1, Th2, Th17 and Treg subsets, suggesting Th1/Th2/Th17/Treg decision-making to be a two-step process. Our results reveal multistable dynamics and establish the stable co-existence of hybrid cell-states, offering a potential explanation for simultaneous differentiation of multipotent naïve CD4+ T cells.

## Introduction

Multistability – the co-existence of more than one steady state/phenotype – is a hallmark of gene regulatory networks (GRNs) driving cellular differentiation and reprogramming. It is the defining trait of a switch that allows the ability to achieve multiple states, without altering internal genetic content [1,2]. It has been observed in diverse biological contexts – lactose utilization in *E. coli* [3], flower morphogenesis in plants [4], multisite phosphorylation [5], haematopoiesis [6], and cancer cell plasticity [7,8]. Thus, decoding the emergent dynamics of underlying regulatory networks is crucial for mapping the cell-fate trajectories and for designing synthetic multistable circuits [9].

Toggle switch – a mutually inhibitory feedback loop between two nodes – is a network motif that enables multistability. A toggle switch between two master regulators A and B often leads to co-existence of two mutually exclusive cell-states – (low A, high B) and (high A, low B) [10,11]. These states correspond to differentiated phenotypes that a common progenitor cell can give rise to; for instance, a toggle switch between PU.1 and GATA1 enabling the common myeloid progenitor to differentiate into a myeloid (high PU.1, low GATA1) or erythroid (low PU.1, high GATA1) cell-state. Intermediate cell-states – (medium A, medium B) – corresponding to hybrid phenotypes have also been observed in scenarios of A and/or B self-activating themselves directly or indirectly [2,8,10]. However, the emergent dynamics of GRNs involved in differentiation of a common progenitor into more than three phenotypes have not yet been as well-studied.

CD4+ T-cells offer an intriguing model system to investigate multistable dynamics with plasticity seen among multiple CD4+ T-cell subsets both *in vitro* and *in vivo* – Th1, Th2, Th9, Th17 and Treg. Specific cytokines can polarize naïve CD4+ T-cells towards these different subsets. Each T-cell subset has unique cytokine production and immune function profile, and retains the capacity to reprogram to other cell-states when exposed to different cytokine environments [12]. The lineage-specifying transcription factors corresponding to Th1, Th2 and Th17 – T-bet, GATA3 and RORγT – have been shown to repress each other, thus forming a toggle triad [13]. Our previous work showed that a toggle triad between A, B and C can enable the co-existence of three differentiated states – (high A, low B, low C), (low A, high B, low C) and (low A, low B, high C) and switching among them. Moreover, three hybrid or ‘double-positive’ states – (high A, high B, low C), (low A, high B, high C) and (high A, low B, high C) were also observed, albeit at a lower frequency than the differentiated ones. These results could explain the experimentally observed phenotypic switching among Th1 (high T-bet, low GATA3, low RORγT), Th2 (low T-bet, high GATA3, low RORγT), Th17 (low T-bet, low GATA3, high RORγT), and the hybrid Th1/Th2, Th1/Th17 and Th2/Th17 states [14,15].

Besides the Th1/Th2/Th17 toggle triad, CD4+ T-cells could also differentiate to regulatory T cells (Treg) that are immunosuppressive in nature, with FOXP3 acting as the master regulator [12]. FOXP3 can inhibit and is inhibited by T-bet, GATA3 and RORγT directly or indirectly [16–21]. Also, double positive cells co-expressing T-bet and FOXP3, GATA3 and FOXP3, and RORγT and GATA3 have been reported, suggesting the presence of hybrid Th1/Treg, Th2/Treg and Th17/Treg states [12]. However, it remains unclear whether a four-node mutually repressive network among T-bet, GATA3, RORγT and FOXP3 is sufficient to explain the co-existence of these 10 states – four single-positive ones (Th1, Th2, Th17 and Treg) and six double-positive ones (Th1/Th2, Th1/Th17, Th2/Th17, Th1/Treg, Th2/Treg and Th17/Treg) and switching among them.

Here, we investigated the emergent dynamics of a toggle tetrahedron – four nodes (A, B, C and D) repressing each other, by simulating a set of coupled differential equations over a parameter ensemble. We show that this network predominantly allows six double-positive states: (high A, high B, low C, low D), (high A, low B, high C, low D), (high A, low B, high C, low D), (low A, high B, high C, low D), (low A, high B, low C, high D) and (low, low B, high C, high D). The presence of single-positive states, on the other hand, is much less prevalent. We further demonstrate switching among these states, and identify the network design principles enabling the co-existence of these states. Our results suggest that differentiation of a progenitor cell into four distinct single-positive states is a two-step process: first, it acquires one of the six double-positive (hybrid) states, following which one of the two lineages is chosen. They also offer a mechanistic explanation for how a ‘toggle tetrahedron’ among T-bet, GATA3, RORγT and FOXP3 allows for the stable existence of multiple intermediate T-cell subsets.

## Results

### Toggle tetrahedron enables six predominant ‘double-positive’ states

Previous reports have identified pairwise mutual inhibition between lineage-specifying transcription factors of Th1, Th2, Th17 and Treg: T-bet, GATA3, RORγT and FOXP3 respectively [13,16–21]. We had earlier investigated the emergent dynamics of a toggle triad, reflecting interlinked toggle switches among T-bet, GATA3 and RORγT. Here, we incorporate the toggle switch that FOXP3 forms with each of these three factors, thus forming a toggle tetrahedron (TTr) – a four-node mutually repressive network (**Fig 1A**). To further verify the antagonism that FOXP3 has with T-bet, GATA3 and RORγT, we analysed RNA-sequencing data for distinct CD4+ T-cell subsets sorted from peripheral blood of healthy donors (GSE135390) including Th1, Th2, Th17, Treg and hybrid Th1/Th17 ones [22]. We quantified the enrichment of previously identified Th1, Th2, Th17 and Treg specific gene lists [23,24] in these subsets. We observed that Th1 gene signature was relatively enriched in Th1 and hybrid Th1/Th17 subsets compared to Th2, Th17 and Treg (**Fig 1B, i**). Similarly, Th2 gene signature was enriched in Th2 cells, Treg signature in Treg cells and Th17 signature in Th17 and hybrid Th1/Th17 cells (**Fig 1B, ii-iv**). These trends suggest the enrichment of Th1, Th2, Th17 and Treg signatures in corresponding cell types.

**Figure 1:**
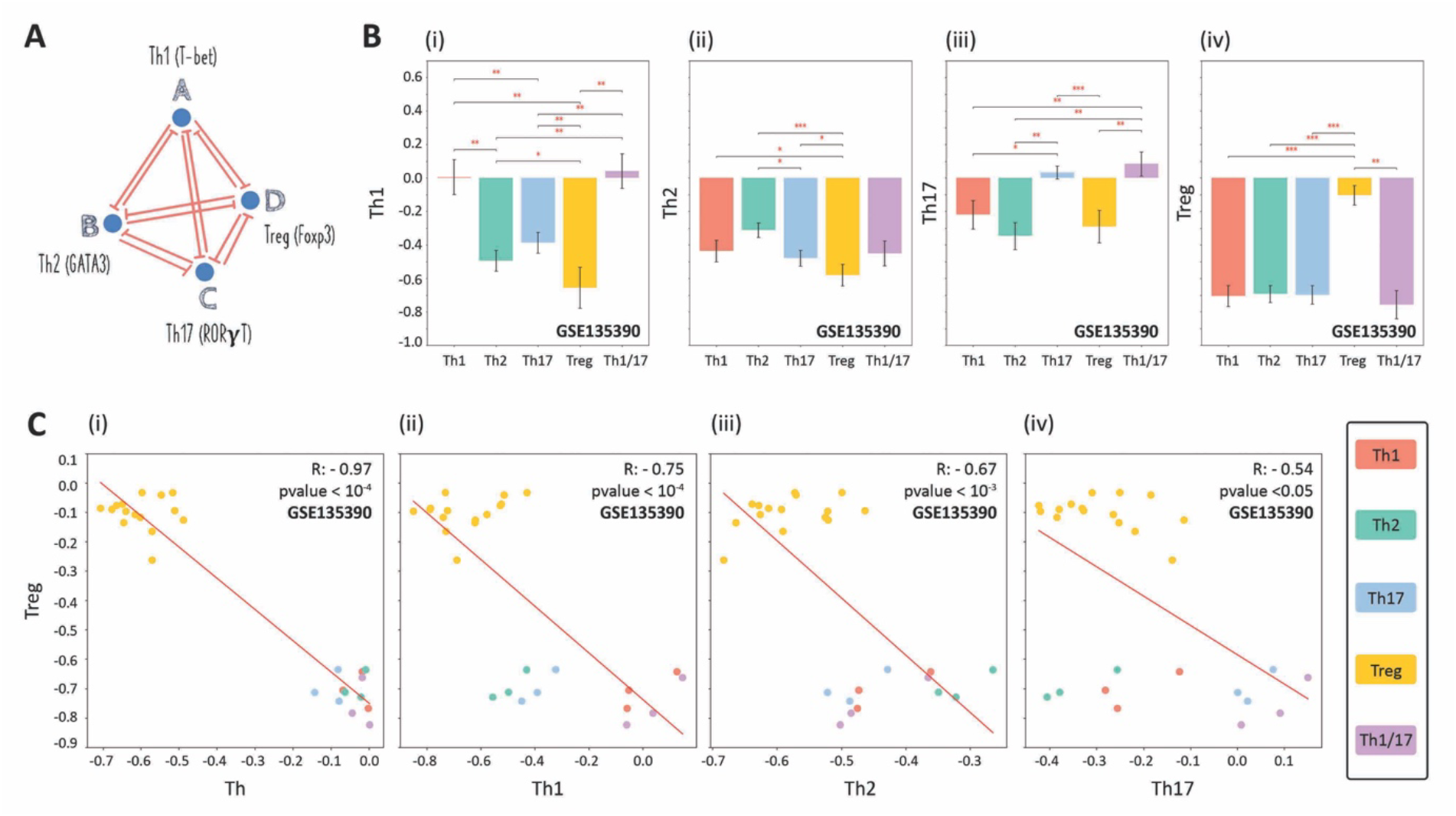
Transcriptomic analysis showing enrichment of Th1, Th2, Th17 and Treg signatures corresponding to specific cell type. **A)** Schematic showing the regulatory network among the regulators of Th1 (T-bet), Th2 (GATA3), Th17 (RORγt) and Treg (Foxp3), forming a toggle tetrahedron. Quantification of difference in levels of i) Th1, ii) Th2, iii) Th17 and iv) Treg gene signature enrichment scores across Th1, Th2, Th17, Treg and hybrid Th1/17 cells (GSE135390). **C)** i) Scatterplot showing different cell types on the Th-Treg gene signature enrichment score plane. ii-iv) Same as i) but for Th1-Treg plane, Th2-Treg place and Th17-Treg plane (GSE135390). Pearson’s correlation coefficient values are shown. *: p-value < 0.05, **: p-value < 0.01; ***: p-value <0.0001 for Students’ two-tailed t-test.

Further, we project these scores on a scatterplot that highlights that Treg specific gene signature enrichment is negatively correlated individually with the enrichment of Th1 (r = - 0.75, p <0.0001), Th2 (r = - 0.67, p < 0.001) and Th17 (r = - 0.54, p< 0.05) specific gene signatures (**Fig 1C, ii-iv**) as well as with a common Th specific gene signature seen in Th1, Th2 and Th17 cells (r = - 0.97, p < 0.0001) (**Fig 1C, i**). Together, these results establish the mutual antagonism that the lineage-specific transcription factor for Treg (FOXP3) has with those of Th1, Th2 and Th17 - T-bet, GATA3 and RORγT respectively. Integrating these trends with a toggle triad observed among T-bet, GATA3 and RORγT [14], we establish the formation of a toggle tetrahedron among these four factors.

We next investigated the dynamical properties of a toggle tetrahedron (between A, B, C and D) using a computational tool, Randomized Circuit Perturbation (RACIPE) analysis [25]. The input to RACIPE is a network topology – a list of activating and inhibiting interactions among the different nodes. RACIPE converts the topology into a set of coupled ODEs (ordinary differential equations) that reflect the set of interactions in that network topology. It then samples 10,000 unique sets of kinetic parameters over a biologically relevant range of values, and generates an ensemble of mathematical models, each with a unique combination of parameter set values. For each such set, RACIPE randomly samples multiple initial conditions for each node in the network, simulates the dynamics of the network topology and reports different possible steady-state values for each node. It should be noted that for some parameter sets, depending on initial conditions, the system may converge to more than one steady state, showcasing multistability for those specific sets. Thus, each kinetic model simulated via RACIPE represents a distinct parameter combination, denoting the inherent cell-to-cell variability in biochemical reaction rates. An ensemble of such models can, therefore, represent the behaviour of a cell population.

Here, each kinetic model is a set of four coupled ODEs. Each ODE tracks the levels of a node engaged in a toggle tetrahedron: A, B, C and D. Next, we characterized the different steady states/ phenotypes enabled by toggle tetrahedron over all parameter sets as identified by RACIPE. Among the 10,000 parameter sets generated for the toggle tetrahedron, the network enabled about 17% monostable cases, 34% bistable cases, 26% tristable cases, 12% tetrastable cases, 4% penta-stable cases and 7% cases of more than 5 co-existing states (**Fig 2A**), thus highlighting the underlying multistable behaviour for this network topology. To identify which specific states are enabled by the network over all multistable sets, we normalized the expression levels of the four nodes A, B, C and D for all solutions corresponding to up to 5 co-existing states (93% of solutions) and plotted them as a heatmap (**Fig 2B**). The heatmap shows the predominance of six states where two nodes were expressed higher relative to the other two (‘double-positive’ states) – {high A, high B, low C, low D}, {low A, low B, high C, high D}, {high A, low B, low C, high D}, {low A, high B, high C, low D}, {high A, low B, high C, low D} and {low A, high B, low C, high D} – represented as {ABcd}, {abCD}, {AbcD}, {aBCd}, {AbCd} and {aBcD} respectively, hereafter. The dominance of ‘double positive’ states was maintained upon varying the number of parameter sets chosen (10^5^, instead of 10^4^) and initial conditions per parameter set (10^4^, instead of 10^3^) (**Fig S1**). Because each node in a TTr is capable of exhibiting ‘high’ or ‘low’ levels, the network can have a total of 16 (= 2^4^) states. We quantified the frequencies of these 16 states over all parameter sets and observed a clear dominance of the six possible ‘double-positive’ states followed by the existence of ‘triple-positive’ and ‘single-positive’ states (**Fig 2C**). The six ‘double-positive’ states ({ABcd}, {abCD}, {AbcD}, {aBCd}, {AbCd} and {aBcD}) each accounted for about 15% of all the states occurring. The four ‘triple-positive’ states ({ABCd}, {AbCD}, {ABcD} and {aBCD}) each accounted for about 2% of all states occurring. The ‘all-high’ and ‘all-low’ states ({ABCD} and {abcd}) and the four ‘single-positive’ states ({Abcd}, {aBcd}, {abCd} and {abcD}) were the least prevalent. Among only the parameter sets enabling monostability, a similar trend repeats with each of the six ‘double-positive’ states each accounting between 14-16% of the cases while each of the four ‘single-positive’ and four ‘triple-positive’ states each accounting for about 1.5% of the cases (**Table S1**), reflecting the symmetry of a toggle tetrahedron. We have considered only the ‘double-positive’ states for further analysis since the rest together do not have more than 10% frequency.

**Figure 2:**
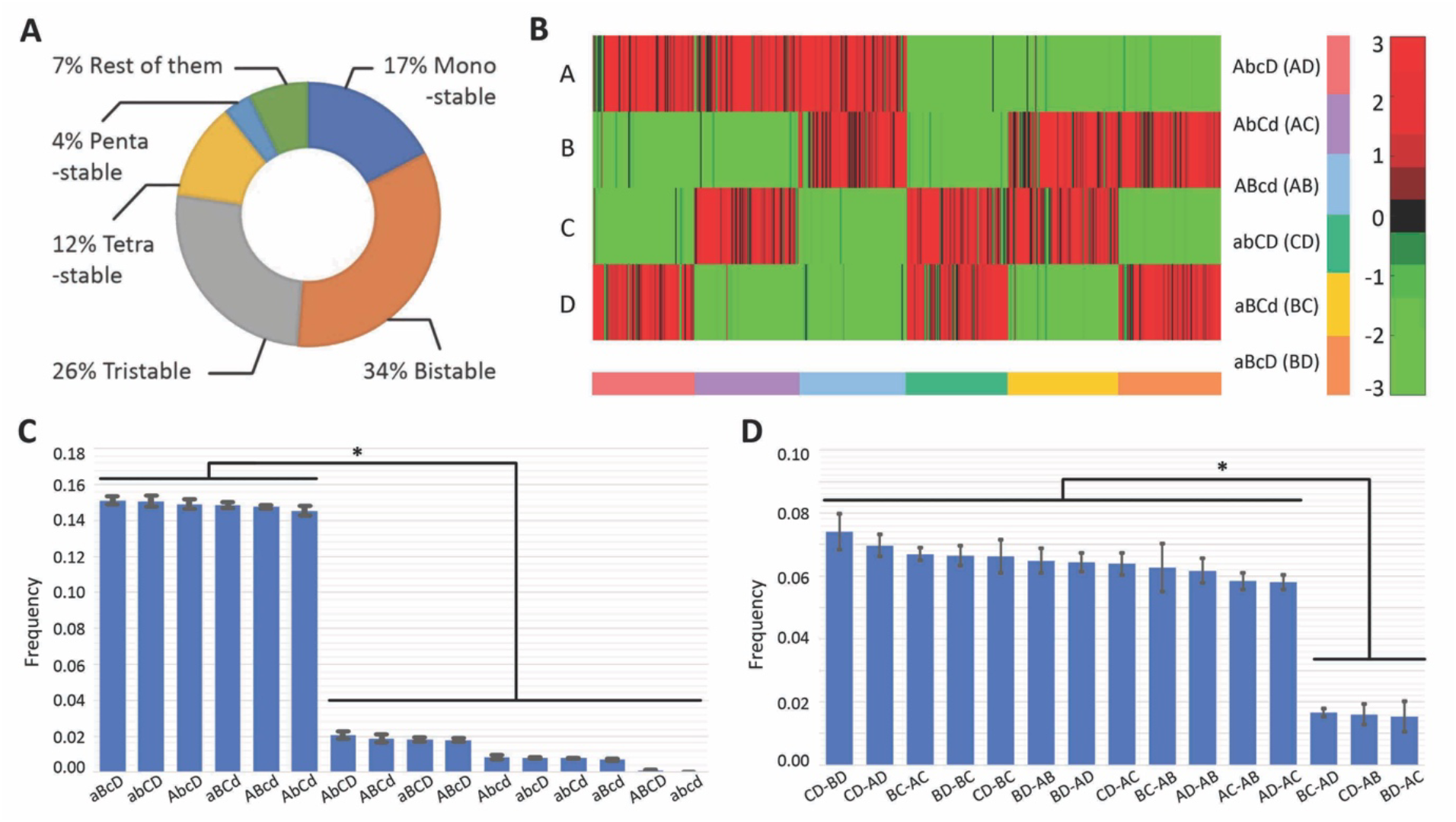
Characterization of phenotypes enabled by a toggle tetrahedron. **A)** Frequency of monostable, bistable, tristable, tetrastable, pentastable solutions and more than 5 co-existing states in the toggle tetrahedron, shown as a pie chart. **B)** Heatmap showing the solutions obtained via RACIPE and nomenclature shows nodes with high (low) expression level in an uppercase (lowercase) fashion. Frequency distribution of states (all solutions taken together) with the most frequent ones highlighted being – {aBcD}, {abCD}, {AbcD}, {aBCd}, {ABcd} and {AbCd}. **D)** Frequency distribution of 15 possible bistable combinations. For the panels C & D, RACIPE data was collected from three independent runs. * shows p < 0.05 for Students’ two-tailed t-test.

Next, we characterized the combinations of steady states given by bistable parameter sets. A bistable parameter set would give rise to two stable steady states. Thus, a total of 120 (=^16^C2) possible bistable combinations are possible, but only 15 combinations had a frequency of more than 1% as a fraction of all bistable combinations, and the remaining 105 (= 120 – 15) combinations only accounted for 19% frequency (**Table S1**). Thus, we ignored these 105 combinations for further analysis and focused on those 15 ones also. Not surprisingly, these 15 (= ^6^C2) combinations are sets of any two ‘double-positive’ states, given their predominance in monostable parameter sets as well as all parameter sets together. Intriguingly, out of the 15 possible combinations of bistable states - 3 states which are a combination of mirror states (i.e., when the bistable state is – {ABcd, abCD} / {AbCd, aBcD} / {AbcD, aBCd}) were found to have significantly lower frequency (together accounted for about 4.5% of the parameter sets – each accounting for about 1.5%) than the other 12 states which consisted of combination of non-mirror states (together accounted for 84% of the parameter sets – each accounting for about 7%) (**Fig 2D**). Put together, our results suggest that the toggle tetrahedron network topology allows for the co-existence of ‘double-positive’ states/ phenotypes where the expression level of two of the regulators is higher relative to the other two.

In addition, we use the complementary tool Dynamic Signatures Generated by Regulatory Networks (DSGRN) [26,27] to analyse the behaviour of TTr across the set of all parameters, as we did earlier for a toggle switch and toggle triad [28]. DSGRN analyses behaviour of ODE models with piecewise constant nonlinearities, which result from taking a limit of the Hill function nonlinearities used by RACIPE as the Hill coefficient tends to infinity. This approximation enables DSGRN to divide high-dimensional parameter space into a finite number of regions defined by explicit inequalities among parameters and identify the corresponding stable steady states/phenotypes associated to all real-valued parameters within that region. By ignoring the value of Hill coefficient, RACIPE parameters may be directly assigned to a DSGRN parameter region, permitting an analytical comparison of phenotype predictions of the TTr model to RACIPE output. Since DSGRN computes steady states by combinatorial methods and does not use ODE simulations, the computational time using DSGRN is several orders of magnitude smaller compared to that of RACIPE. However, the TTr provides a challenge to the combinatorial methodology of DSGRN, as the number of parameter regions of the TTr is over 27 trillion. Instead of an exhaustive computation of steady states for all these regions, we introduce two approaches. First, we explore a very small (6561 regions out of 27 trillion) but well-studied subset of the parameter regions that we term Strict monotone Boolean (SB) parameters. This approach corresponds to a choice of monotone Boolean function at each vertex of TTr and studying the number of steady states for the corresponding Boolean dynamics. Second, we perform stochastic sampling of DSGRN parameter regions. Finally, we compare the frequencies of different types of steady states predicted by DSGRN SB parameters and by DSGRN stochastic parameter sampling to RACIPE stochastic samples.

We first compared the predictions of DSGRN SB parameter subset to the predictions obtained by stochastic sampling of DSGRN parameter space. We notice substantial differences in predictions. DSGRN SB parameters indicate a much greater frequency of double-positive states than stochastic sampling (**Fig 3A**, top). Additionally, DSGRN SB parameters predict that 12 of the 15 possible bistable states exhibit much higher frequencies than the remaining 3 bistable states, while DSGRN stochastic sampling only shows a small difference between these two subsets of bistable states (**Fig 3A**, bottom). Next, we compared the all-state frequency predictions of DSGRN to those of RACIPE (top row) and similarly the frequency predictions of bistable states (bottom row). This comparison was done to both DSGRN stochastic sampling (**Fig 3B**) and to DSGRN SB parameters (**Fig 3C**). We observe that low frequency predictions in RACIPE are correlated with low frequency predictions in the DSGRN methods, and similarly for higher frequency predictions. However, the best affine fit between DSGRN and RACIPE predictions has a linear coefficient other than 1 (blue regression lines vs black line in **Figs 3B-3C**). The DSGRN SB parameter predictions have a linear coefficient closer to 1 than the stochastic samples, and in this sense may indicate closer relationship between the predictions of RACIPE and DSGRN SB than between RACIPE and DSGRN stochastic sampling. Lastly, and for completeness, we show the best affine fit between DSGRN stochastic sampling and DSGRN SB parameters for all states (**Fig 3D** top) and bistable states (**Fig 3D** bottom). As expected from the comparisons to RACIPE, the low and high frequencies are well correlated, but the linear coefficient differs from 1.

**Figure 3:**
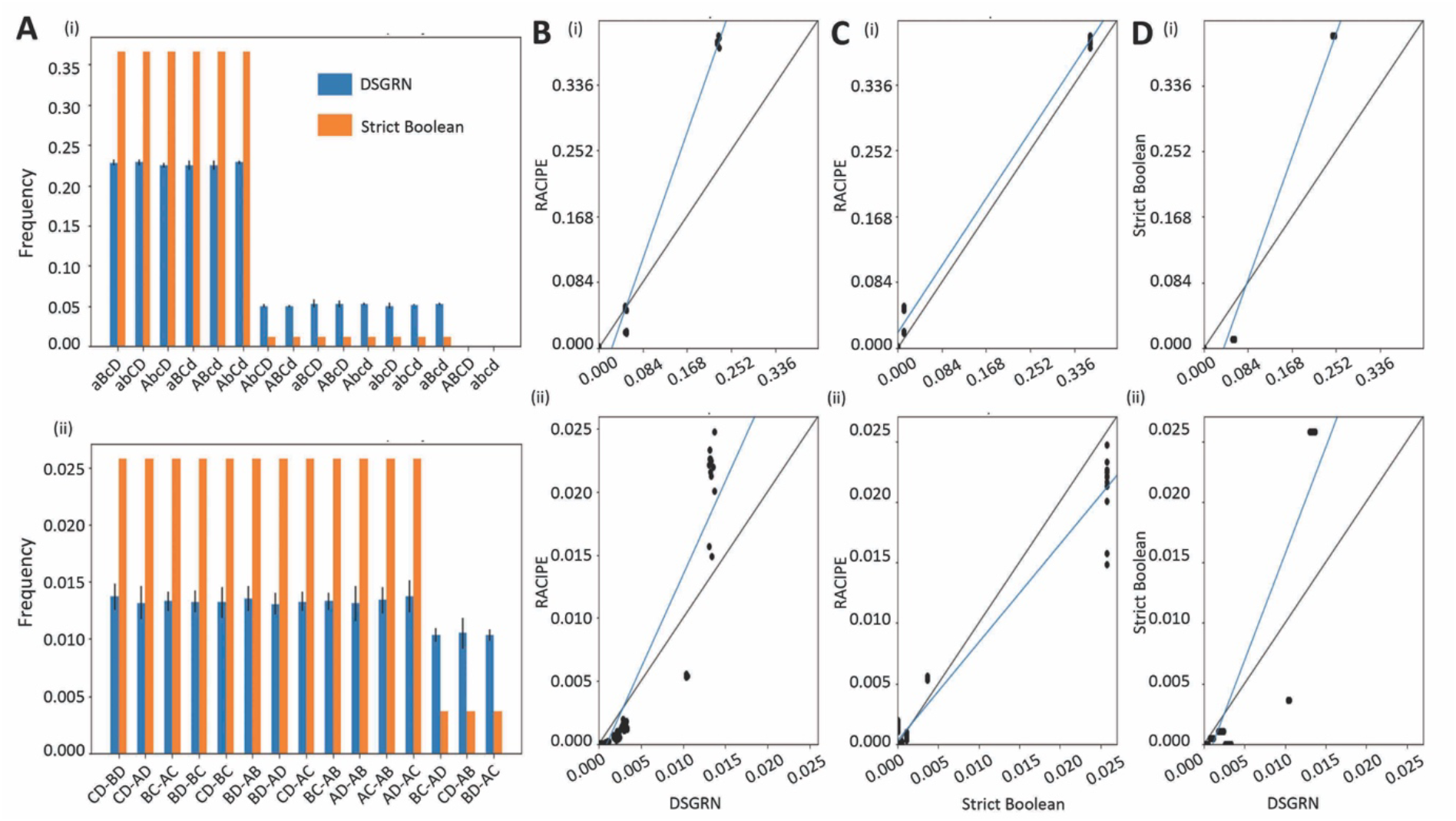
State-space analysis of toggle tetrahedron using DSGRN. **A)** Frequency of all states (top) and frequency of bistable states (bottom) for the DSGRN randomly sampled parameters (blue) and all 6561 strict monotone Boolean parameters (orange). Note that four sets of 10,000 parameters were sampled, the black line on the blue bars indicate the standard deviation in frequency between these sets. **B)** Simple linear regression [29] was used to show the relationship between RACIPE and sampled DSGRN frequencies (black dots) for all states (top) and bistability (bottom). The line of best fit is indicated by the blue line, while the black line represents the line y=x. **C)** Same as (B) for RACIPE and strict Boolean frequencies. (D) Same as (B) with strict Boolean and sampled DSGRN frequencies. Notice that when the blue line slope and intercept are close to the black line, as we see in (C), indicates that the frequencies between the pairs are very similar

We show that the steady states of SB parameters where each monotone Boolean function is nondegenerate, i.e. non-constant and where each input is able to affect the output, can be analytically determined (**SI Section 3**). We use this analysis to confirm the numerical results (**Fig 2C, Fig 3A**) that the double positive equilibria occur more frequently than other types of equilibria. We further analyse other symmetric tetrahedron networks where number of positive in-edges at each node is either 1, 2 or 3. Our analysis shows that in each of these cases within the ensemble of all compatible non-degenerate monotone Boolean functions, the frequency of double positive states is higher than the frequency of other states (**SI Section 3**).

### Dynamical traits of double-positive states enabled by toggle tetrahedron

To further characterize the parametric space corresponding to co-existence of the double-positive states, we performed bifurcation analysis of multiple parameter sets enabling non-mirror bistable states identified by RACIPE. For a representative parameter set enabling the bistable state – {ABcd-AbcD}, where the expression level of nodes B and D should switch, we chose the degradation rate of B (kB) as the bifurcating parameter (**Fig 4A,i**). We observed that at high levels of kB, the system loses bistability and becomes monostable as the state {ABcd} no longer exists. Conversely, at very low levels of kB, only the state {ABcd} exists and the system loses bistability. In the bistable regime (kB ranging between 0.1 and 0.8, thus spanning almost an order of magnitude), the expression levels of B and D switch between two states – one is relatively high and the other low. The nodes A and C also exhibit two states, but the levels of A are high in both the states, and those of C are relatively too low to distinguish between them. For the kB value obtained from the RACIPE generated parameter set, we performed stochastic simulations to validate the switching between states and observed the expression levels of B and D (**Fig 4B,i**). We see that the levels of B and D are mutually exclusive i.e., when B is relatively high, D is low and *vice versa*, as expected from the bistable state where B and D switch between states.

**Figure 4:**
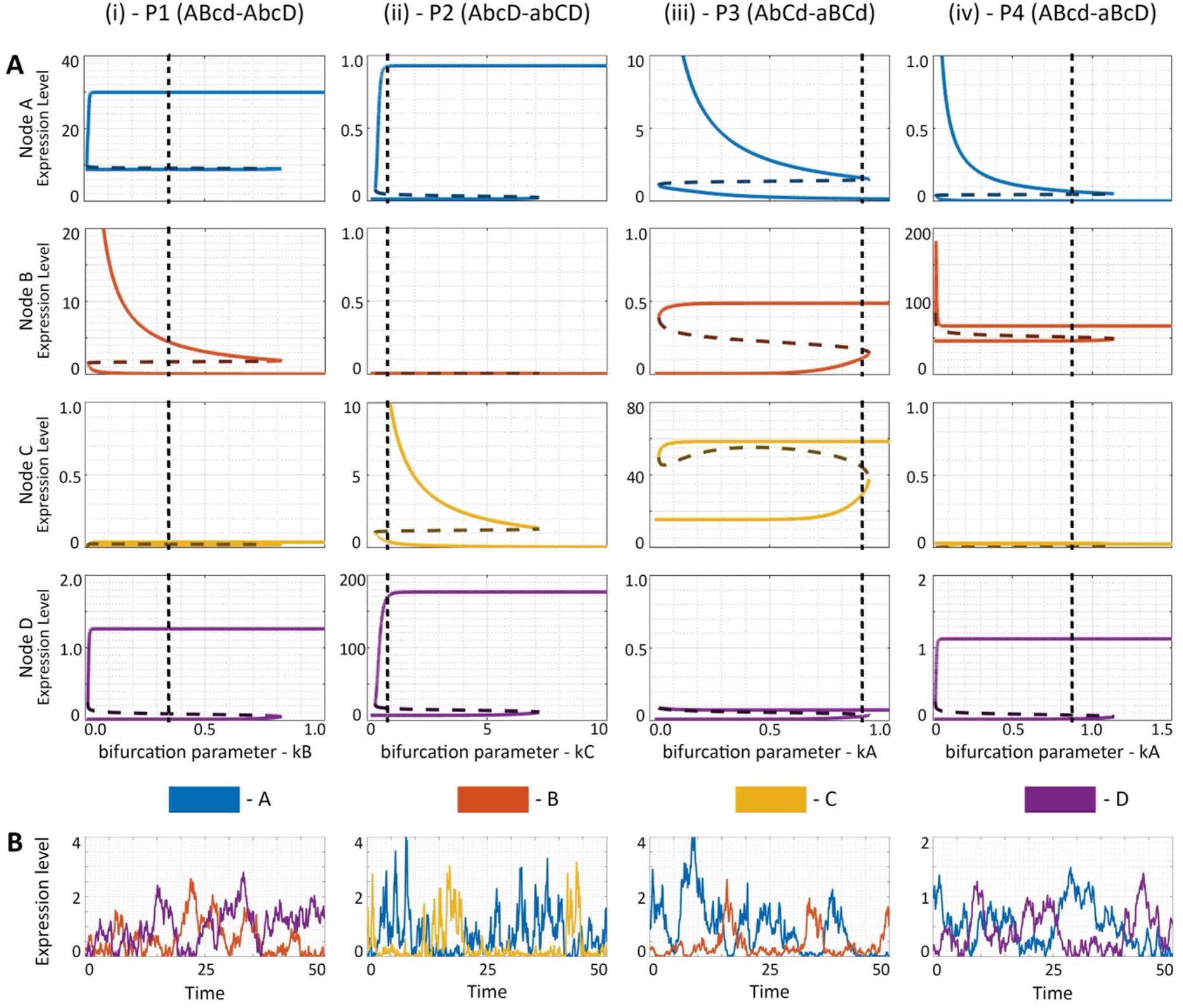
Bifurcation analysis of representative parameter sets corresponding to the non-mirror bistable states. **A)** Bifurcation diagrams of four representative cases (each column denotes a different parameter set – P1-P4) with each row showing the expression levels of the four nodes A, B, C and D. Dotted vertical line in each case marks the RACIPE parameter value of the corresponding bifurcation parameter. **B)** Stochastic simulations showing switching between the bistable states (only the two nodes switching between ‘high’ and ‘low’ are shown, the nodes always expressed ‘high’ or always ‘low’ are not shown). Parameter sets P1,P2, P3 and P4 are provided in Table S2.

We performed the same analysis for three more representative parameter sets enabling the bistable states – {AbcD, abCD}, {AbCd, aBCd} and {ABcd, aBcD}, and observed similar behaviours though the bifurcating parameters were degradation rates of different nodes (kC, kA and kA respectively) (**Fig 4A,ii-iv**). In each of the cases, we see that the expression level of the node which is always ‘high’ switches between two levels but both these levels are relatively ‘high’. Stochastic simulations showed possible switching between the two bistable states enabled by the respective parameter set (**Fig 4B, ii-iv**). Put together, these results highlight that co-existence of ‘double positive’ states is a robust feature of TTr and this feature can be disrupted by altering the degradation rate of nodes in the TTr network (i.e. the half-life of one of the TFs engaged in a TTr is drastically manipulated).

### Design principles of multistability enabled by toggle tetrahedron

A toggle tetrahedron can exhibit monostable (17% of parameter sets) or multistable (83% of parameter sets) dynamics. Thus, we investigated various parameter sets from the RACIPE analysis to deduce a relationship between the parameter sets and the different states they converged to. We hypothesized that the relative strength of transcriptional inhibition among the different nodes controls the different stable steady states observed. To quantify the strength of inhibition from one regulator onto the other, we used a metric called link strength X, where X(AB) represents the value of inhibition from regulator A to B, in accordance with the formulation of RACIPE framework [14].

In RACIPE formalism, the strength of the inhibitory interaction is defined by a shifted Hill function consisting of three parameters – *n* (Hill coefficient corresponding to cooperativity), λ (fold-change) and T (half-maximal concentration or threshold). For an inhibitory link, the higher the value of n, the faster is the increase in strength of inhibition with changing concentration of the source node, or in other words, the steeper the increase in strength of inhibition. For inhibitory links, λ value ranges from 0.01 (strong repression) to 1 (no effect). Thus, the smaller the value of λ, the stronger the inhibition. Similarly, the smaller the value of T, the lower is the concentration of source regulator needed for the inhibition to be active. Thus, we defined the link strength metric, X= n/(λ *T), such that the higher the value of X, the stronger is the strength of inhibition.

For the parameter sets that enable the monostable ‘single-positive’ state {abcD} or {low A, low B, low C, high D}, we hypothesize that the inhibitions of A, B and C by D are relatively stronger than inhibition of D by A, B and C. To test the hypothesis, we first shortlisted the parameter sets that enable the particular state (here, monostable {abcD}), and then calculated the strength of inhibition between each pair of nodes for each parameter set. For every parameter set, and for each pair of nodes, among the two inhibitory links, we identified the dominant one and thus quantified the frequency with which one node inhibits the other one more strongly compared to *vice versa*. We found that for approximately 70% of the parameter sets, the conditions X(DA) > X(AD), X(DB) > X(BD) and X(DC) > X(CD) were true, i.e. inhibitions originating from the node D were stronger than their opposing counterpart in a mutually inhibitory feedback loop (**Fig 5A**). On the other hand, no such skew was observed for mutual inhibition among pairs of nodes whose levels were low, i.e. between A and B, between B and C and between A and C, i.e. in approximately 50% of parameter sets, X (AB) > X (BA), and in the remaining 50%, X (BA) > X (AB) (**Fig 5A**). Results for other monostable ‘single-positive’ states show similar trends (**Fig S2A-C**). For the case of monostable ‘double-positive’ state {abCD} or {low A, low B, high C, high D}, we hypothesized that the inhibition of A and B by C and D is stronger than that of C and D by A and B. We found our hypothesis to be true in 70% of the parameter sets corresponding to this state, i.e. the conditions X(CA) > X(AC), X(DB) > X(BD), X(CB) > X (BC) and X (DA) > X (AD). However, the two remaining pairwise mutual inhibitions – one between A and B, and the other between C and D – did not show any such skew (**Fig 5B**). Consistent trends were observed for other monostable ‘double positive’ states (**Fig S2D-F, S3B-F**). Similarly, for a mono-stable ‘triple positive’ state {aBCD} or {low A, high B, high C, high D}, we noticed that in about 70% of corresponding parameter sets, the inhibition of A by B, C and D was higher than inhibition of B, C and D by A, i.e. X (BA) > X (AB), X (CA) > X (AC) and X (DA) > X (AD). However, no such skew was noted for pairwise mutual inhibition among B, C and D (**Fig S3A**). These trends highlight the parametric conditions under which TTr topology does not allow for multistable behaviour and instead converges to one of the different possible monostable scenarios. Theoretical analysis of nondegenerate monotone Boolean functions also confirms the RACIPE results that for any pair of genes the strength of repressive connection from highly expressed gene is larger than that from a weakly expressed gene (**SI section 4**).

**Figure 5:**
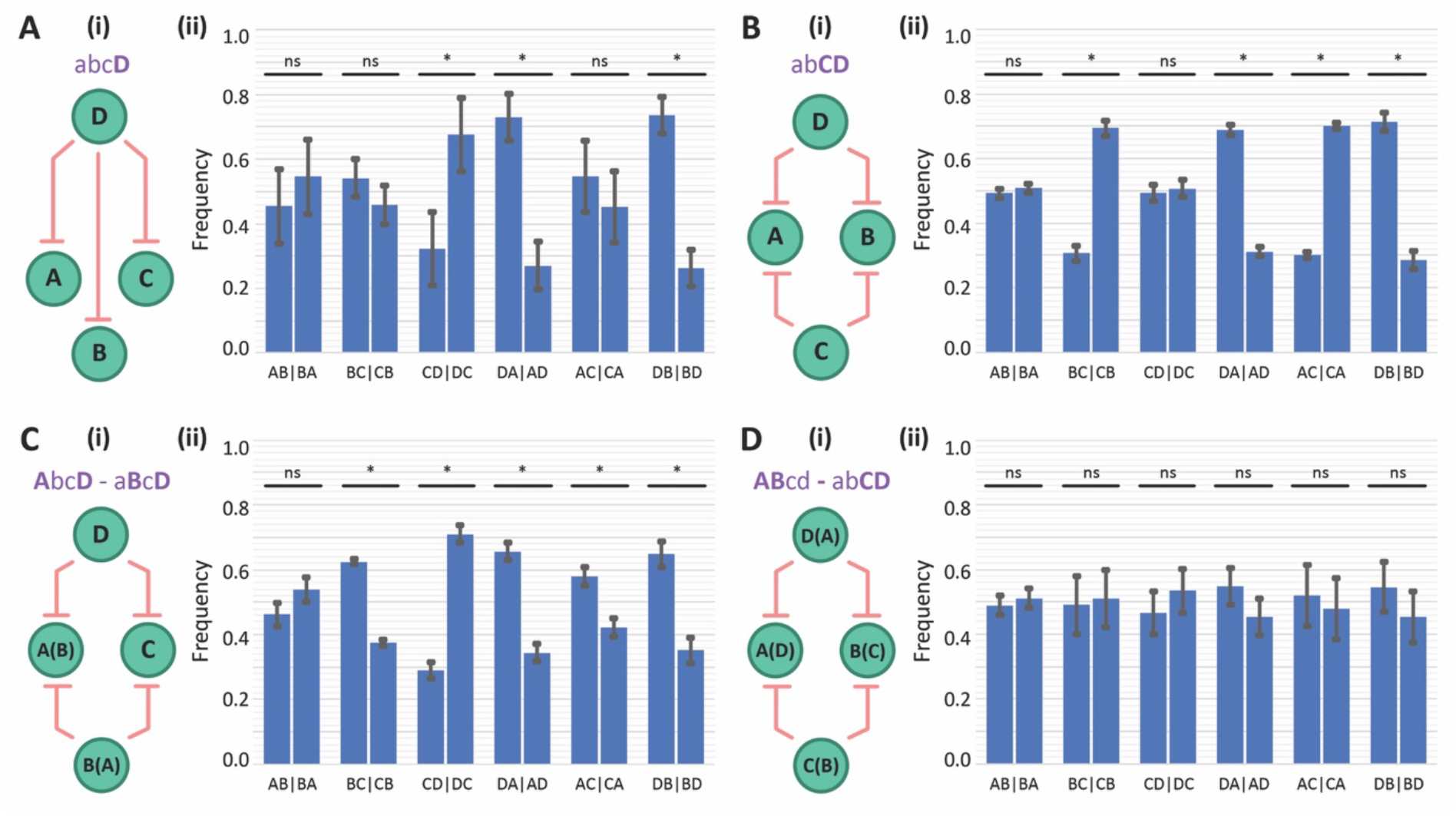
Link Strength Analysis of monostable and bistable states for toggle tetrahedron. **A)** i) Schematic showing the links that are expected to be stronger than their counterparts for the state {abcD} ii) Frequency of dominance of all six pairs of mutually inhibitory links between any two nodes in a toggle tetrahedron, for all parameter sets corresponding to the case of monostable {abcD}. **B)** Same as A) but for the monostable state – {abCD}. **C)** Same as A) but for the non-mirror bistable state combination – {AbcD, aBcD}. **D)** Same as A) but for the mirror bistable state combination – {ABcd, abCD}. In D) and panels, the nodes that switch between the two states are written as X(Y) where X and Y can switch.

Next, we performed the link strength analysis for parameter sets enabling bistability. In the case of bistable states, we have two predominant combinations among the ‘double-positive’ states – a pair of mirror states and a pair of non-mirror states. Mirror states refer to the combination of states where all four nodes switch their levels between the two states, say {abCD, ABcd}. Non-mirror states refer to the combination of states where two nodes do not switch their levels, say {aBcD, AbcD}, where A and B switch their levels between the two states, but C and D do not.

For parameter sets enabling the non-mirror state combination of {aBcD, AbcD}, we hypothesized the following: a) inhibition of A, B and C by D is overall stronger than inhibition of D by A, B and C; ii) inhibition of C by A, B and D is overall stronger than that inhibition of A, B and D by C; iii) for mutual inhibition between A and B, both the links are equally likely to be stronger than the other. An analysis of corresponding parameter sets reveals our hypothesis to be true: i) X(DC) > X(CD) in approximately 70% of parameter sets, X(DA) > X(AD) and X(DB) > X(BD) in approximately 65% of the parameter sets, ii) X(AC) > X(CA) and X(BC) > X(CB) in approximately 60% of parameter sets, and X(DC) > X(CD) in about 70% of parameter sets, and iii) X(AB) > X(BA) in about 50% of the parameter sets (**Fig 5C**). In the case of parameter sets enabling bistability with the pair of mirror states – {abCD, ABcd}, we hypothesized that for mutual inhibition between every pair of nodes, each inhibition is equally likely to be stronger than the other. The results showed that the for the two inhibitions between any pair of nodes, one is stronger than the other in about 50% of the parameter sets (**Fig 5D**). Similar trends were observed for the other bistable state combinations (**Fig S4-S6**). Put together, these results point towards patterns in a high-dimensional parametric space that allows a toggle tetrahedron to enable specific combinations of states or dynamical behaviours.

### Uniqueness of the dynamical traits of toggle tetrahedron

T-bet, GATA3, RORγT and FOXP3 – similar to many master regulators – are known to self-activate directly and/or indirectly [14,30,31]. Thus, we investigated the dynamics of TTr when all 4 nodes can either self-activate (TTr+SA) or self-inhibit (TTr+SI). We observed that compared to TTr, for TTr+SA, multistability was enhanced, i.e. a higher number of parameter sets enabling bistability, tristability, tetrastability and pentastability, consistent with previous observations of multistability being associated with total number of positive feedback loops in a given network [32]. However, for TTr+SI, the number of parameter sets enabling multistability decreased and those corresponding to monostability increased (**Fig S19**). Irrespective of these changes, the ‘double-positive’ states were still the most predominant ones in case of both TTr+SA and TTr+SI (**Fig 6A**), showcasing that the salient features of a TTr remain unchanged upon adding self-regulations.

**Figure 6:**
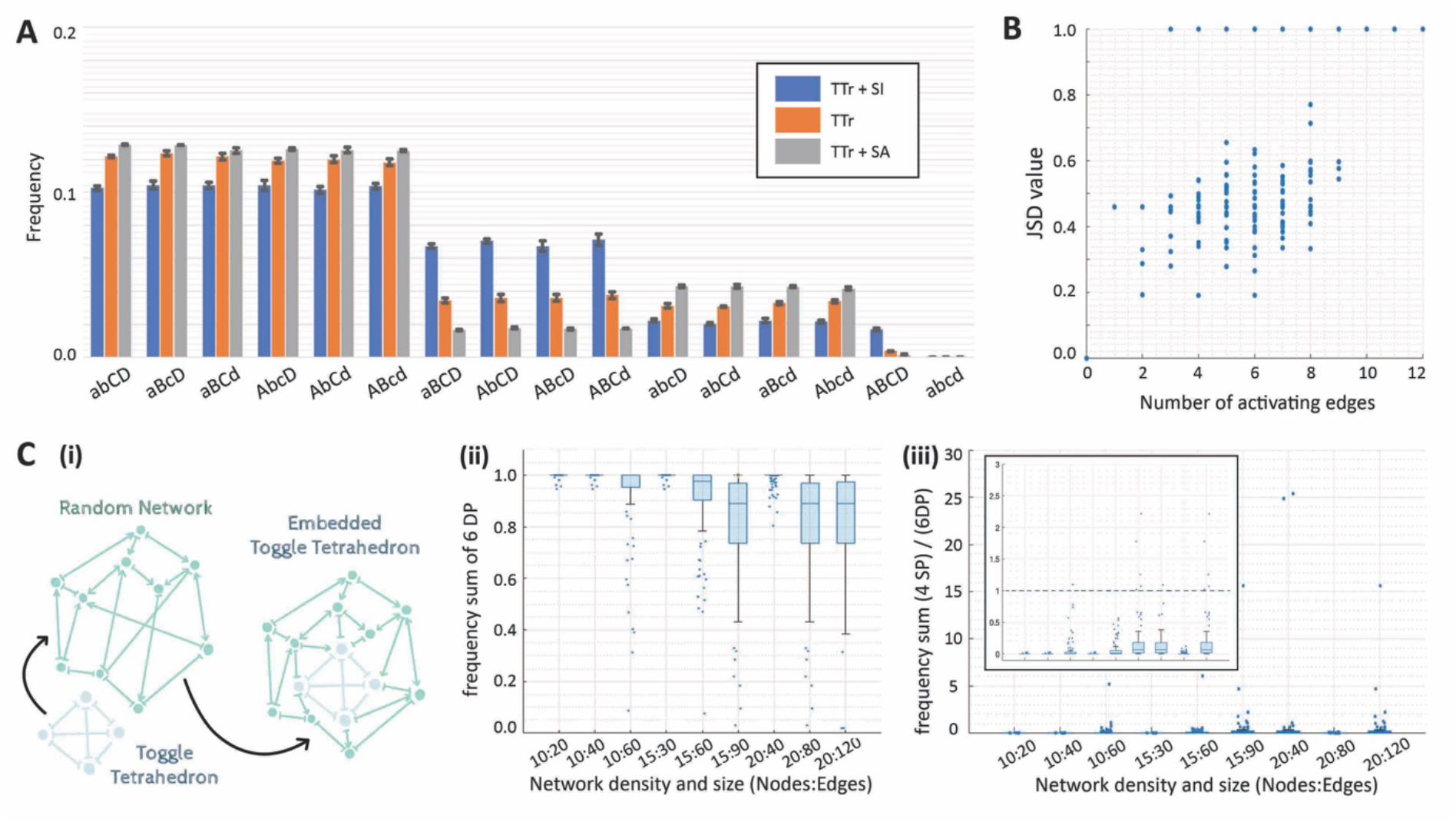
Features unique to the toggle tetrahedron topology. **A)** Frequency of monostable, bistable, tristable and other multistable states for TTr, TTr+SI and TTr+SA. **B)** Scatterplot showing the JSD value between the frequency distribution graphs of the reference network (TTr) and the network specified by the number of inhibiting edges swapped (on x-axis) when all steady states are taken together. **C)** i) Schematic representing how a TTr network is embedded in a larger random network. ii) Sum of the frequencies of all the six ‘double-positive’ states for Boolean simulations of networks of varying sizes and densities. iii) Fraction of sum of frequencies of all the four ‘single-positive’ states (4 SP) over the sum of frequencies of all the six ‘double-positive’ states (6 DP) for Boolean simulations of networks of varying sizes and densities. Inset shows a zoomed in version with the y-axis range – (0,3).

We next asked how unique is the predominance of ‘double-positive’ states to a TTr topology. To answer this question, we calculated the steady-state distributions of all possible fully connected four-node networks, i.e. networks in which each node is connected to at least one of the other 3 nodes. No self-regulatory edge was allowed in this ensemble of networks; given that each of the 12 edges connecting any two nodes can be activatory or inhibitory, we have a large possibility space of network topologies. We shortlisted them, using the NetworkX library in Python [33], to ensure that each network is unique, i.e. no network is repeated in the network topology space. We found a total of 218 possible networks including the TTr. Next, we compared the behaviour of the 217 networks, one at a time, with the TTr as the reference by measuring how different or similar the frequency distribution of the 16 possible states. To minimize computational costs of simulating 217 networks, these simulations were done using discrete Boolean modeling for general asynchronous update scheme and simple majority update rule, instead of RACIPE. Previous simulations for other network topologies showed remarkable consistency in steady-state distributions obtained via RACIPE and Boolean modeling [8,32,34], thus we chose this modeling strategy. We used JSD (Jensen-Shannon Divergence) as our metric to compare corresponding steady-state distributions. JSD values lie between 0 to 1; the higher the value, the more different the two distributions are. We noticed that as we transition from TTr to other four node networks in which one or more of inhibitions between two nodes is swapped with an activatory edges, the higher the number of such swaps, the higher the JSD (**Fig 6B**). Intriguingly, for as less as three swaps, we begin to see networks with JSD = 1, i.e. completely non-overlapping steady-state distributions, when compared with the TTr. JSD was found to be 0 only when there were no swaps. Put together, this result shows that the features observed for TTr are unique to its network topology and increasingly differ with increasing number of swaps of inhibitions by activations in the network topology.

Regulatory networks such as a TTr do not operate in isolation, but are often embedded in larger networks. Thus, we quantified the behaviour of TTr when embedded in external networks of varying sizes and densities (**Fig 6C, i**). We chose these external networks of 3 sizes (10 nodes, 15 nodes, 20 nodes) and 3 densities (no. of edges = 2* no. of nodes, no. of edges = 4* no. of nodes, no. of edges = 6* no. of nodes). For a given size and density, we chose 100 unique random networks in which we embedded the TTr. Thus, in total, we simulated 900 (= 3*3*100) networks, and used the abovementioned Boolean modeling strategy to minimize computational costs. For each of these 900 networks, we measured the net frequency of the six ‘double-positive’ phenotypes, and plotted the distribution of these frequencies for the set of 100 external networks corresponding to a fixed size and density. We noticed that as the size and more importantly, the density of the external network increased, the sum of frequencies of the ‘double positive’ states decreased (**Fig 6C, ii**). However, the ‘double positive’ states continued to be quite dominant, and in most cases, the six ‘double positive’ states remain much more frequent than the four ‘single positive’ ones (**Fig 6C,iii**), reflecting the functional resilience of a TTr topology, when embedded in external networks. This behaviour is reminiscent of the resilience of toggle switch and toggle triad when embedded in similar external networks, where the ‘single-positive’ states are the most dominant states (**Fig S7**) [35].

## Discussion

Multistability is a hallmark of diverse cell-fate decision regulatory networks [2], and synthetic multi-stable circuits are being increasingly integrated in *E. coli*, yeast and mammalian cells [9,36–38]. It is not observed only at an intracellular level, but also in spatial tissue-level patterning through the emergent dynamics of cell-cell communication networks such as Notch-Delta-Jagged signaling [39,40]. Thus, decoding the dynamical traits of multistable networks is critical to better understand cellular development and reprogramming and has attracted extensive theoretical attention too [41]. Most deterministic or stochastic computational models of multistable networks have assessed the dynamics of a mutually inhibitory loop between two transcription factors (TFs), or a TF and micro-RNA, where the TFs may self-activate [2,42–45]. Such toggle switches are extensively observed at cell-fate bifurcation points in developmental decision-making [10], where a common progenitor can give rise to two mutually exclusive phenotypes. Consistently, a toggle switch between nodes A and B can allow for co-existence of (high A, low B), (low A, high B) and (medium A, medium B) states. Here, we have investigated the dynamics of four lineage-specific TFs mutually repressing one another, as observed in CD4+ T-cell differentiation [12], thus forming a toggle tetrahedron. Our results suggest that a toggle tetrahedron allows for the predominant existence of six ‘double-positive’ states where two of the four lineage-specific TFs have relatively higher expression levels as compared to the other two. This behaviour is fundamentally different from a toggle switch or a toggle triad, where ‘single-positive’ (only one of the lineage-specific TF is at relatively high levels) states dominate. Extending these results to understand the differentiation of naïve CD4+ T cells into Th1, Th2, Th17 and Treg subsets, we can indicate that this differentiation is a two-step process, where a multipotent cell first acquires one of the hybrid or ‘double-positive’ state (step 1), and further differentiate into one of the ‘single-positive’ states after the hybrid/progenitor cell switches to one of the two possible phenotypes (step 2) (**Fig 7**). Moreover, our results suggest that all the six ‘double-positive’ states are not merely intermediaries, but can be relatively stable phenotypes, albeit with possibly higher plasticity compared to differentiated Th1, Th2, Th17 and Treg subsets. Stable hybrid Th1/Th2 (T-bet+ GATA3+) [46], hybrid Th17/Treg (RORγT+ FOXP3+) [47] and hybrid Th1/Treg (T-bet+ FOXP3+) [21] phenotypes have been reported experimentally, thus validating our model predictions. Further time-course high-throughput analysis of CD4+ T-cell differentiation [15,22,48] into multiple (> 2) subsets simultaneously would help in testing our prediction about the two-step differentiation.

**Figure 7:**
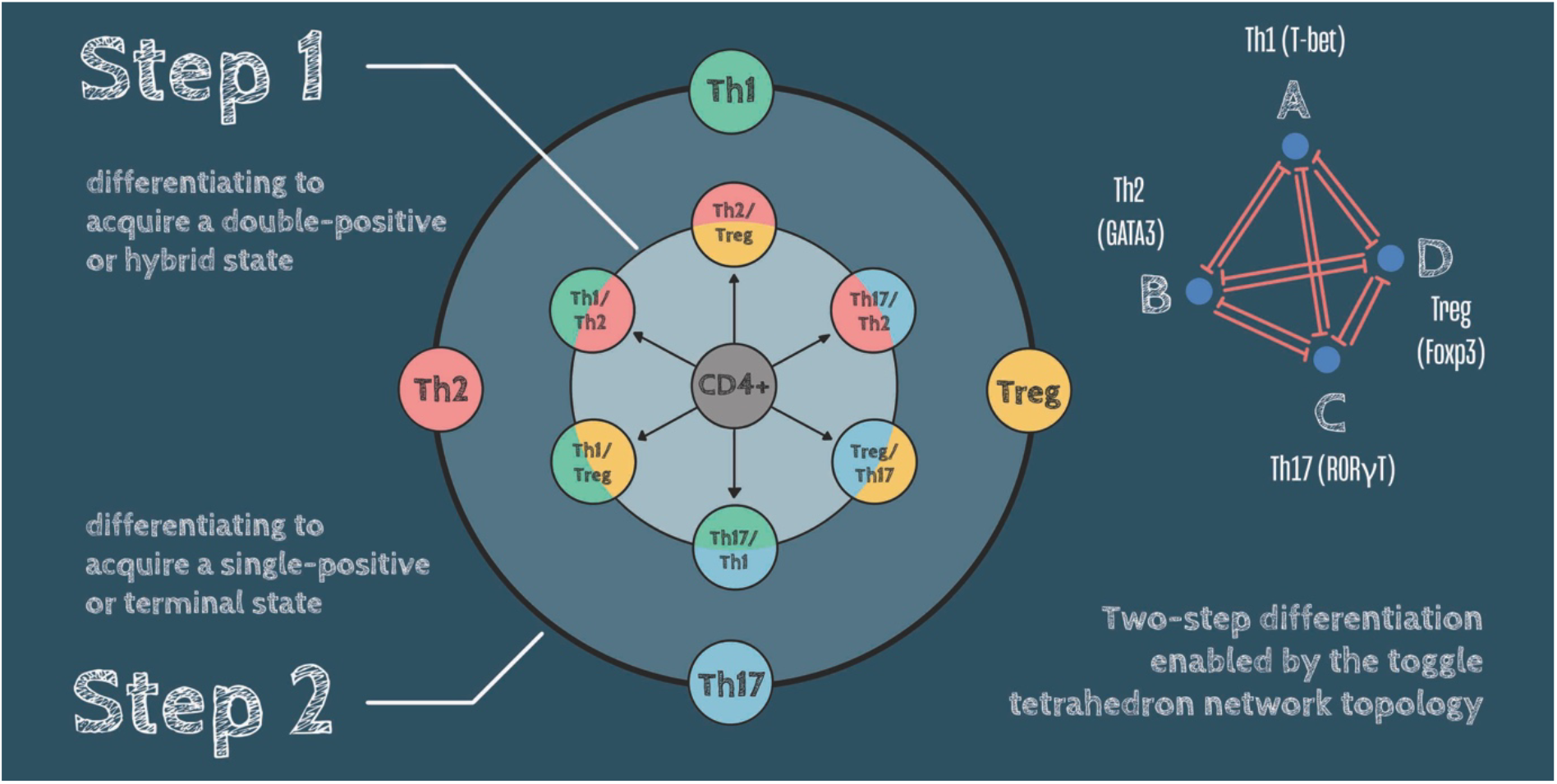
Schematic representing the two-step differentiation enabled by the toggle tetrahedron. (left) CD4+ T-cells can acquire hybrid or ‘double-positive’ states and differentiate into a single-positive state, as enabled by dynamics of toggle tetrahedron among T-bet, GATA3, RORγt and Foxp3 (right).

Previous computational models of CD4+ T-cell differentiation have often focused on the dynamics under a specific parameter set, and have reported the presence of hybrid phenotypes [19,49,50]. Our analysis builds on those efforts and quantifies the frequency of different states across a wide parameter set ensemble, thereby demonstrating the salient features of the TTr network topology. Importantly, this predominance of ‘double positive’ state in a TTr is not seen in any other four-node network, and is retained even when it is embedded in larger networks, thus offering insights into the resilience of TTr network dynamics. Future computational models of CD4+ T-cell differentiation should incorporate additional lineage-specific TFs such as BCL-6 and Blimp-1 (that regulate the T follicular helper cell differentiation) and their interactions with T-bet, GATA3 and FOXP3 [51–55]. Increasing size of the network presents challenges to analytical and numerical exploration of their dynamics. Larger networks come with larger number of parameters and RACIPE sampling thus covers less of the parameter space. The number of parameter regions (i.e. DSGRN parameters) examined by DSGRN methodology also grows exponentially with the size of the network. However we observed that within the small subset of 6561 SB parameters out of 27 trillion of all parameters for TTr there is surprisingly high correlation between the results from RACIPE sampling, SB parameter and sampling of all DSGRN parameters exhibited. We are currently seeking explanation of this phenomena, since examining only SB parameters would allow analysis of much larger networks.

Overall, our results offer novel insights into TTr dynamics and propose a step-wise decision-making for a common progenitor cell capable of giving rise to more than two differentiated phenotypes.

## Methods

### Randomized Circuit Perturbation (RACIPE) analysis

RACIPE [25] is a computational tool used to investigate the dynamical behaviour exhibited by a specific network topology. It attempts to quantify all possible steady-state behaviours the network topology can show over an ensemble of parameter sets, instead of a specific parameter set. Thus, it samples parameters over a biologically relevant range to generated multiple parameter sets. A large number of parameter sets is likely to ensure that all diverse behaviours in parametric space are accounted for. Analysing these results provides an understanding of the different states enabled by the topology in specific parametric spaces and an idea of the frequencies with these states occur.

The formulation of one-way interaction between any two nodes of the network topology (say an inhibition from node B onto node A) is given by the following equation:

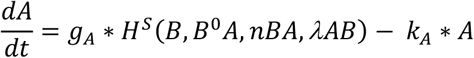

where *g*_*A*_ and *k*_*A*_ are intrinsic production and degradation rates of the node A respectively and the Hill function *H*^*s*^(*B, B*^*0*^*A, nBA,λAB*) represents the interaction (here inhibition) of the node B onto node A. The first term on the RHS represents the net production rate of node A while the second term represents the degradation of node A (here, a first order degradation is considered). The Hill interaction consists of a combined form of negative and positive Hill functions and is referred to as a shifted Hill equation. The Hill function is used to represent the activation or inhibition between two nodes because of the use of biochemical rate equation formulation of gene expression.

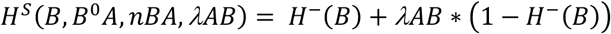

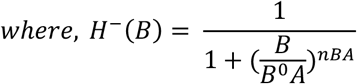

In case of the interaction being an activation, the hill function is further divided by the fold-change parameter corresponding to the respective interaction i.e., in case of an activation from node B to node A, the hill function would be, *H*^*s*^(*B, B*^*0*^*A, nBA, λAB*)/ *λAB*.

The default range of values for Hill coefficient in RACIPE is [1, 6], but we chose the range of [6,10] for a TTr, because it allows for a bimodal distribution for the node expression levels, to segregate ‘high’ and ‘low’ states (**Fig S1A**). We used the default values of number of parameter sets (=10000) and number of initial conditions per parameter set (=1000) for our simulations, although similar behaviour was observed when taking a larger number of parameter sets and/or initial conditions (**Fig S1B**). The threshold values are calculated such that for each parameter set, every interaction has a 50% chance of being active/inactive. For every parameter set, classification of its monostability, bistability or other multistability is done after simulating it for the 1000 initial conditions and finding the number of stable steady states obtained.

### Finding SF value for TTr embedded in larger networks

We embedded the TTr motif in randomly generated networks of specific sizes and densities i.e., specific number of nodes and edges in the network. For a given pair of values for number of nodes and edges, we generated 100 networks randomly without overlap and for each network we ran Boolean simulations following asynchronous update rule as specified by (). We then calculated the sum of frequencies for the six ‘double-positive’ states (SF) enabled by the TTr motif.

### NetworkX – for finding unique four node topologies

The constraints for us to find all possible network topologies is that they have to have four nodes and each node is connected to every other node via an activation or inhibition. We first generated all possible topologies with every edge having a possibility of an activation or inhibition. To simplify the code, we did not consider notations of activations and inhibitions but just marked the presence of an edge to be an activation and considered its absence to be an inhibition. We then generated all possible combinations of edges between nodes. For every classification based on number of edges present, we compared the network topologies within each class using the `nx.algorithms.is_isomorphic` function from NetworkX [33]. The `nx.algorithms.is_isomorphic` function utilizes an implementation of the VF2++ algorithm [56] for Graph Isomorphism testing to shortlist the unique network topologies in our case. Finally, after obtaining the unique topologies, we converted them to complete directed topologies (i.e., marked activations and inhibitions again). We exported these directed network topologies in the ‘.topo’ file format for further RACIPE analysis.

### Dynamic Signatures Generated by Regulatory Networks (DSGRN) analysis

DSGRN is a computational tool devised to perform an exhaustive computation of coarse dynamics (e.g., steady states) across all possible parameter values of a switching system [28]. The switching system for the TTr is given by four equations, one for each node. The equation for node A is given; the equations for the other nodes are analogous.

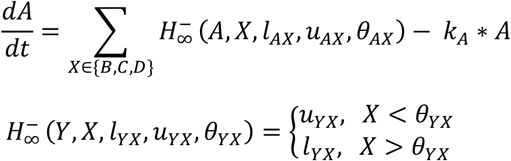

Notice that, 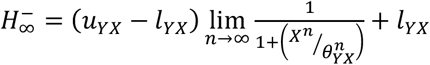 which permits a mapping from RACIPE parameters to DSGRN parameter regions (see [28] for details). We refer to *l*_*YX*_, *u*_*YX*_, *θ*_*YX*_ as the lower, upper, and threshold values of the edge *X → Y*.

The parameter regions defined by DSGRN are given by collections of inequalities derived from the topology of the TTr, one for each of A, B, C, and D. Each node has three input edges, and therefore has eight potential input values (example shown for node A in the first column of **Table 1**). These input values to A are interleaved with the thresholds *θ*_*XA*_ of A’s output edges to form a DSGRN parameter region for the A node. An example set of DSGRN parameter inequalities for node A is given in the 2^nd^-4^th^ columns of **Table 1**, where the red text highlights less than (<) relationships.

**Table 1.** Column 1: Input values to node A. Columns 2-4: A choice of input value relationships to output thresholds. As discussed in the text, this table represents an example of two DSGRN parameters for node A.

| Input value | $\theta_{BA}$ | $\theta_{CA}$ | $\theta_{DA}$ |
| --- | --- | --- | --- |
| $I1 = l_{AB}l_{AC}l_{AD}$ | $I1 < \theta_{BA}$ | $I1 < \theta_{CA}$ | $I1 < \theta_{DA}$ |
| $I2 = u_{AB}l_{AC}l_{AD}$ | $I2 > \theta_{BA}$ | $I2 > \theta_{CA}$ | $I2 < \theta_{DA}$ |
| $I3 = l_{AB}u_{AC}l_{AD}$ | $I3 > \theta_{BA}$ | $I3 > \theta_{CA}$ | $I3 < \theta_{DA}$ |
| $I4 = l_{AB}l_{AC}u_{AD}$ | $I4 > \theta_{BA}$ | $I4 > \theta_{CA}$ | $I4 > \theta_{DA}$ |
| $I5 = u_{AB}u_{AC}l_{AD}$ | $I5 > \theta_{BA}$ | $I5 > \theta_{CA}$ | $I5 < \theta_{DA}$ |
| $I6 = u_{AB}l_{AC}u_{AD}$ | $I6 > \theta_{BA}$ | $I6 > \theta_{CA}$ | $I6 > \theta_{DA}$ |
| $I7 = l_{AB}u_{AC}u_{AD}$ | $I7 > \theta_{BA}$ | $I7 > \theta_{CA}$ | $I7 > \theta_{DA}$ |
| $I8 = u_{AB}u_{AC}u_{AD}$ | $I8 > \theta_{BA}$ | $I8 > \theta_{CA}$ | $I8 > \theta_{DA}$ |

The table above implicitly defines *θ*_*BA*_ *< θ*_*DA*_ and *θ*_*CA*_ *< θ*_*DA*_; however, the relationship of *θ*_*BA*_, *θ*_*CA*_ remains an open choice, and therefore the table represents two DSGRN parameter regions. The dynamics and thus the different possible steady-state values for each node are determined by a choice of DSGRN parameter [27]. For the TTr, there are 3 in-edges and 3 out-edges from every node, resulting in 4242 parameter regions for each node. Each node independently may be associated to any one of these inequalities, resulting in 4242^4^ DSGRN parameter regions, an incomputable number. Instead of exhaustive computations, we use both stochastic sampling and a principled restriction of the number of DSGRN parameter regions.

### Stochastic simulations

As previously stated, the number of DSGRN parameter regions for TTr makes exhaustive computing infeasible. We randomly sampled four sets of 10,000 DSGRN parameters, for a total of 40,000 parameters. For each sample group, we used the DSGRN software (https://github.com/marciogameiro/DSGRN) to calculate the discrete dynamics of each parameter, and hence the classification of its monostability, bistability or other multistability.

### Strict monotone Boolean parameters

Modelling using monotone Boolean functions has a long history [57,58]. DSGRN parameters are readily described as multilevel Boolean functions [59], which have a special subset of monotone Boolean functions. We restrict our attention to these DSGRN parameter regions, that we call strict monotone Boolean (SB) parameters. The set of SB parameters are all DSGRN parameter regions (i.e., they respect the partial order in **Table 2**) such that an input value *Ik* is either less than or equal to all the output thresholds of node A. An example SB parameter is given in **Table 2**, where again less than relationships are highlighted in red. Notice that every row (i.e., each input value) is either all red or all black, this is the condition to be an SB parameter. We note that there are additional algebraic constraints on DSGRN parameters. Specifically, the input values (i.e., the lower and upper values) into a node are evaluated as a sum of products, see [27] for more details. In a SB parameter there are no input values between any two thresholds, and therefore the thresholds may occur in any of the six possible orders. Therefore, a SB parameter table always represents six DSGRN parameter regions. Given the small size of this collection of parameters (6561) compared to all parameters, the high correlation between the results from RACIPE sampling, SB parameter and sampling of all DSGRN parameters exhibited in **Fig 3B-3D** is surprising. We are currently seeking explanation of this phenomena.

**Table 2.** Column 1: The input value to node A. Columns 2-4: A choice of input value relationships to output thresholds. As discussed in the text, this table represents an example of six DSGRN SB parameters for node A.

| Input values | $\theta_{BA}$ | $\theta_{CA}$ | $\theta_{DA}$ |
| --- | --- | --- | --- |
| $I1 = l_{AB}l_{AC}l_{AD}$ | $I1 < \theta_{BA}$ | $I1 < \theta_{CA}$ | $I1 < \theta_{DA}$ |
| $I2 = u_{AB}l_{AC}l_{AD}$ | $I2 < \theta_{BA}$ | $I2 < \theta_{CA}$ | $I2 < \theta_{DA}$ |
| $I3 = l_{AB}u_{AC}l_{AD}$ | $I3 < \theta_{BA}$ | $I3 < \theta_{CA}$ | $I3 < \theta_{DA}$ |
| $I4 = l_{AB}l_{AC}u_{AD}$ | $I4 > \theta_{BA}$ | $I4 > \theta_{CA}$ | $I4 > \theta_{DA}$ |
| $I5 = u_{AB}u_{AC}l_{AD}$ | $I5 < \theta_{BA}$ | $I5 < \theta_{CA}$ | $I5 < \theta_{DA}$ |
| $I6 = u_{AB}l_{AC}u_{AD}$ | $I6 > \theta_{BA}$ | $I6 > \theta_{CA}$ | $I6 > \theta_{DA}$ |
| $I7 = l_{AB}u_{AC}u_{AD}$ | $I7 > \theta_{BA}$ | $I7 > \theta_{CA}$ | $I7 > \theta_{DA}$ |
| $I8 = u_{AB}u_{AC}u_{AD}$ | $I8 > \theta_{BA}$ | $I8 > \theta_{CA}$ | $I8 > \theta_{DA}$ |

