## Supplementary Text for "Multistability and predominant double-positive states in a four node mutually repressive network: a case study of Th1/Th2/Th17/T-reg differentiation"

#### SUPPLEMENTARY MATERIAL

In this section we develop theory that will allow us to analytically compute number of nondegenerate strict Boolean parameters (SB), explained in the main text, that support double positive equilibria of the type ($ABcd$) vs, single positive ($Abcd$), triple positive ($ABCd$) and all-high ($ABCD$) vs. all low ($abcd$). This will directly validate results in Figure 2C and Figure 3A. Our analysis allows extension of these results from TTr network to other symmetric tetrahedron networks where each node receives 1,2 or 3 positive inputs from the other nodes. In all of these networks the frequency of double positive steady states exceeds those of single, triple and all-high, or all-low steady states.

Finally , in section 4 we analytically support the argument in Fig. 4 about the relative strength of repression in equilibria of type ($abcD$) and ($aBcD$) in TTr network.

##### 1. Symmetric Boolean tetrahedron networks

In this section we will consider symmetric tetrahedron networks with four nodes $A, B, C, D$. We assume that the network is full connected without self-edges and therefore each node receives three inputs from other three nodes. We will consider four tetrahedron networks which are symmetric under full set of permutations of the set $S := \{A, B, C, D\}$ i.e. under an $S_4$ symmetry.

(1) all edges are negative
(2) each node has one positive input edges and two negative input edges. Without loss we will assume that the positive edges connect $A \to B \to C \to D \to A$.
(3) each node has two positive input edges and one negative input edges. Without loss we will assume that the negative edges connect $A \to B \to C \to D \to A$.
(4) all edges are positive.

We associate to each networks (1)-(4) a Boolean network model [5, 6, 7, 4]. Each node $x$ will have two states 0 or 1 representing active vs. inactive state, described the corresponding Boolean variable $X \in \mathbb{B} := \{0, 1\}$. Each state $X$ will be updated by a Boolean function $f_X : \mathbb{B}^3 \to \mathbb{B}$ with $X \in S$. In addition, we assume that each function is monotone [1, 2]

**Definition 1.1.** A function $f : \mathbb{B}^3 \to B$ is non-decreasing (non-increasing) monotone Boolean function (MBF) if

$$f_{|b_i=0} \leq f_{|b_i=1} \qquad f_{|b_i=1} \leq f_{|b_i=0}).$$

for $i = 1, 2, 3$.

The monotonicity of the Boolean function reflects the structure of the regulatory network. When input edge to node $x$ is positive (negative), the monotone Boolean function (MBF) is required to be non-decreasing (non-increasing.

Furthermore, we are only interested in non-degenerate MBFs.

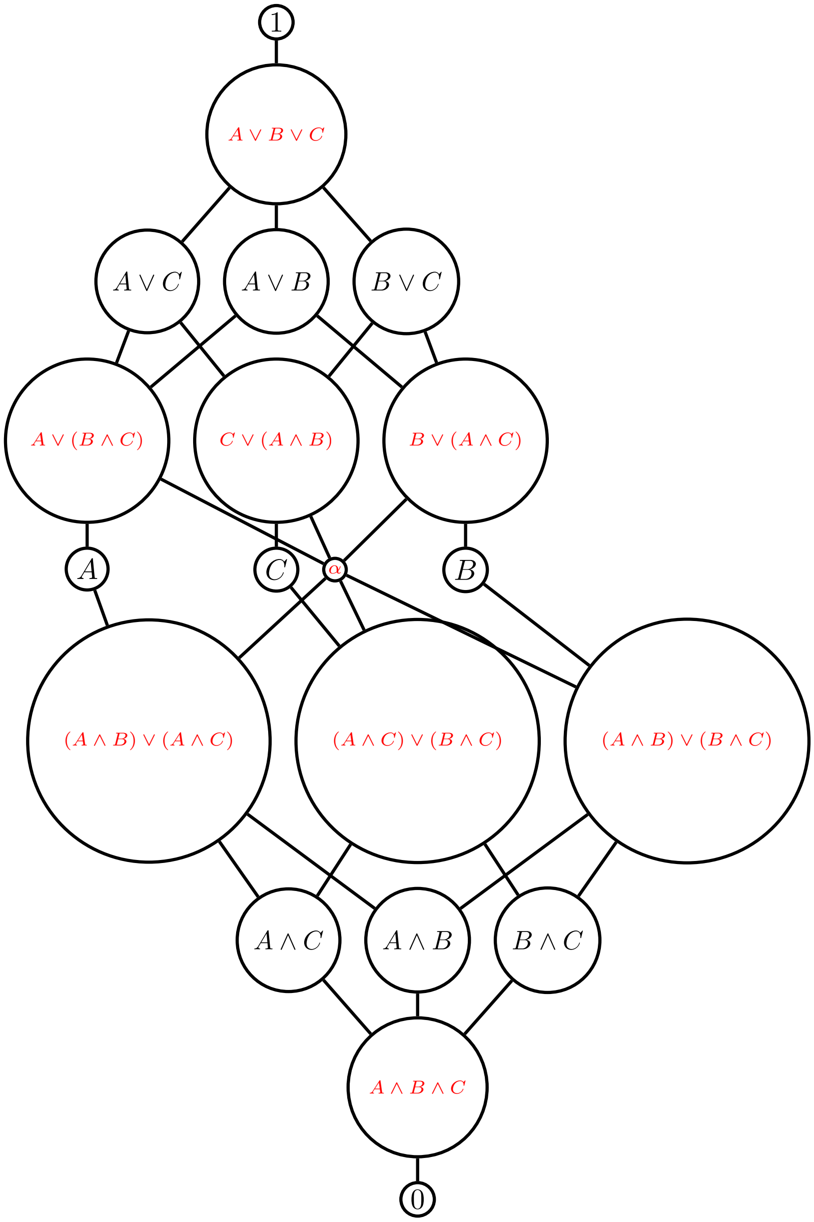


FIGURE 1. Lattice of positive monotone Boolean functions (MBFs) with three inputs $A, B, C$, where $\alpha = (A \wedge B) \vee (A \wedge C) \vee (B \wedge C)$. Note that each functions in each horizontal layer share the size of the truth set, ranging from 0 on the bottom for 0 function to 8 on the top for constant function 1. Red script indicate non-degenerate MBFs.

**Definition 1.2** ([3])**.** A monotone Boolean function $f : \mathbb{B}^k \to \mathbb{B}$ is *non-degenerate* if

(1) $f$ is not constant; and
(2) for every input variable $x$ there exist $b \in \mathbb{B}^k$ such that
$$f_{|b(x=0)} \neq f_{|b(x=1)},$$
where $b(x = a)$ is the word $b$ with entry in the $x$ component equal to $x = a \in \mathbb{B}$.

Lattice of all MBFs with three inputs is in Figure 1 with non-degenerate MBF marked in red [3]. Note that every MBF $f$ with the size of the truth set $\mathbb{T}(f) \in \{1, 3, 5, 7\}$ is non-degenerate, while MBFs with truth set of size $\mathbb{T}(f) \in \{0, 2, 6, 8\}$ are always degenerate.

1.1. **Input ordering.** The lattice in Figure 1 is written in form of non-decreasing Boolean functions; however, the same structure is valid also for inputs that are represented by negative edges. If, for instance, an input edge node $A$ is negative, then the lattice is the same, but every $A$ is replaced by $\neg A$.

For each network we define a partial order on the input set $\mathbb{B}^3$ which is extension of the order induced by each input edge on the set $\mathbb{B}$. In other words $0 < 1$ for a positive edge and $1 < 0$ for a negative edge. Each such ordered Boolean input set $\mathbb{B}^3$ let $M$ is the highest entry and $m$ is the lowest entry. The input state $M$ represents the state where all activating inputs have value 1 and all repressing inputs have value 0. For network (1) the states $M = (000), m = (111)$. On the other hand, for instance for network (3) the top and bottom states will be different for $f_A, f_B, f_C$ and $f_D$. To see this, we note that the inputs to any function $f_X, X \in S := \{A, B, C, D\}$, are the other three variables $S \setminus X$, which we represent in alphabetical order. Therefore for function $f_A$ the inputs are, in order $B, C, D$ while for $f_C$ they are, in order $A, B, D$. With this notation, it is easy to see that for $f_A$ we have $M = (110), m = (001)$ and for $f_C$ we have $M = (101), m = (010)$ which depend on the position of the repressive input edge within the input triple.

#### 2. Non-degenerate MBFs that support equilibria

In this section we will investigate how many non-degenerate MBF $f = (f_A, f_B, f_C, f_D)$ exists that support a particular type of a steady state.

Non-degenerate Boolean functions must satisfy

$$f(M) = 1, \qquad \text{and} \qquad f(m) = 0, \tag{1}$$

as the opposite selection would result in a constant function as a result of monotonicity assumption.

We illustrate our approach on an example. Consider a steady state say $P := (0100)$. Requirement that the function $f$ supports a steady state $P$ gives one constraint on each function $f_X, X \in S$

$$f_A(100) = 0, \quad f_B(000) = 1, \quad f_C(010) = 0, \quad f_D(010) = 0,$$

where the inputs are listed in the alphabetical order as noted above. Each of these conditions may impose additional constraints on other values of the function based on monotonicity. How many of these additional constraints are imposed depends on the value (0 or

1) of the primary constraint and the distance of the input from the maximal input $M$ and the minimal input $m$. Given $M, m \in \mathbb{B}^3$ define two layers in $\mathbb{B}^3$. Let

$$L_m := \{b \in \mathbb{B}^3 \mid |b - m| = 1\}, \qquad L_M := \{b \in \mathbb{B}^3 \mid |b - m| = 2\},$$

where $L_m$ stands for *minimal layer* and $L_M$ for *maximal layer.* As an example, in represilator tetrahedron network where $M = (000)$ and $m = (111)$ we have

$$L_m = \{(110), (101), (011)\}, \qquad L_M = \{(100), (100), (010)\}\}$$

The following Theorem summarizes the number of non-degenerate MBF that satisfy the additional constraint, as a function of which layer the constraint is in, and what is its value. In general, if the constraint is near $M$ and agrees with the value $f(M)$ it is a weaker constraint compared to when the constraint disagrees with the value $f(M)$. Same remark is true for the proximity to $m$.

**Theorem 2.1.** *Assume non-degeneracy condition for MBF $f$*

$$f(m) = 0, \qquad f(M) = 1.$$

*The steady state requirement gives a single constraint for each of $f_A, f_B, f_C, f_D$ at some $b \in \mathbb{B}^3$.*

*There are these possibilities:*

(1) *for $b = M$, we have $f(b) = 1$, or for $b = m$, we have $f(b) = 0$. This gives no additional constraint and there are 9 choices for non-degenerate MBF $f$.*
(2) *for $b \in L_m$, we have $f(b) = 1$, or for $b \in L_M$ we have $f(b) = 0$. This gives rise to 2 non-degenerate MBFs.*
(3) *for $b \in L_M$, we have $f(b) = 1$, or for $b \in L_m$ we have $f(b) = 0$. This gives rise to 7 non-degenerate MBFs.*
(4) *for $b = M$, we have $f(b) = 0$, or for $b = m$, we have $f(b) = 1$. These are incompatible with (1) and gives 0 choices for non-degenerate MBF $f$.*

*Proof.* Before we start the proof we make three observations about the symmetries of the problem, that will allow us to simplify the proof.

First, the set of monotone Boolean functions as well as non-degenerate Boolean functions in Figure 1 is preserved under any permutation of the inputs $A, B, C$. These permutations form a group $S_3$.

Second, the cube of inputs $\mathbb{B}^3$ is also symmetric under the permutations $S_3$. As a consequence, if we prove the statement of the Theorem for, say, constraint $f(001) = 1$, the same result is valid for the constraint $f(\sigma(001)) = 1$ for any $\sigma \in S^3$.

Third, we consider standard dualization operation $D$ (or $\neg$) on Boolean lattice $\mathbb{B}$ that intechanges 0 and 1 and operations $\vee$ and $\wedge$. Note that a non-degenerate monotone Boolean function $f : \mathbb{B}^3 \to \mathbb{B}$ commutes with $D$ i.e. $D \circ f = f \circ D$. As a result the function $g := f \circ D$ satisfying $g(b) = f(D(b))$ has the values that are negation of the values of $f$ on the same input $b$. Therefore $g$ is also a non-degenerate Boolean function that can be obtained from $f$ by replacing inputs $A, B, C$ by $\neg A, \neg B, \neg C$.

Having these three symmetries in mind, we proceed with the proof.

| | |
|---|---|
| 1 | $C$ |
| **2** | $(\mathbf{C} \wedge \mathbf{B}) \vee (\mathbf{C} \wedge \mathbf{A})$ |
| 3 | $B \wedge C$ |
| 4 | $A \wedge C$ |
| **5** | $(\mathbf{A} \wedge \mathbf{B} \wedge \mathbf{C})$ |

| | |
|---|---|
| 1 | $A \vee C$ |
| **2** | $\mathbf{C} \vee (\mathbf{A} \wedge \mathbf{B})$ |
| **3** | $\mathbf{A} \vee (\mathbf{B} \wedge \mathbf{C})$ |
| **4** | $\alpha$ |
| 5 | $C$ |
| 6 | $A$ |
| **7** | $(\mathbf{A} \wedge \mathbf{B}) \vee (\mathbf{B} \wedge \mathbf{C})$ |
| **8** | $(\mathbf{A} \wedge \mathbf{C}) \vee (\mathbf{B} \wedge \mathbf{C})$ |
| **9** | $(\mathbf{A} \wedge \mathbf{C}) \vee (\mathbf{A} \wedge \mathbf{B})$ |
| 10 | $A \wedge B$ |
| 11 | $B \wedge C$ |
| 12 | $A \wedge C$ |
| **13** | $\mathbf{A} \wedge \mathbf{B} \wedge \mathbf{C}$ |

TABLE 1. Logic expression for monotone Boolean functions $f$ in the order of decreasing size of the truth set $\mathbb{T}(f)$ that satisfy constraints in case (2) on the left and case (3) on the right. Bold faced numbers are non-degenerate MBFs.

**Case (1).** This follows directly by counting the number of all non-degenerate MBFs in Figure 1 which is 9.

**Case (2).** Given the permutation symmetry, we consider without loss of generality $m = (000), M = (111)$ and a function $f$ with inputs $A, B, C$ and the constraint

$$f(110) = 0.$$

In addition, we also impose standnig non-degeneracy constraints $f(000) = 0$ and $f(111) = 1$.

Note that since $f(110) = 0$, it follows by monotonicity that $f(b) = 0$ for any $b \prec (110)$. This implies two additional costraints

$$f(100) = 0, \qquad f(010) = 0.$$

This results in five constraints on values of $f$ which only leaves three free values of $f$. If we select all the free values to be 0 we get the function $A \wedge B \wedge C$ in Figure 1; if we replace all free values by 1 we get the function $C$ in Figure 1. Therefore all compatible monotone Boolean functions form a sublattice of the lattice in Figure 1 with the highest element $C$ and the lowest element $A \wedge B \wedge C$. These are listed in Table 1 (left); the 2 nondegenerate MBFs are in bold.

**Case (3).** Given the symmetries, we assume without loss of generality that the function $f$ satisfies constraint

$$f(010) = 0, \text{ together with } f(000) = 0, f(111) = 1. \tag{2}$$

The condition $f(010) = 0$ does not imply any other conditions. There are 13 MBFs listed in Table 1 (right) that satisfy the three conditions in (2). The 7 that are bold are non-degenerate. These functions form a sublattice bounded by the highest function compatible with these constraints which is $A \vee C$ with truth set of size $|\mathbb{T}| = 6$ and the lowest function which is $A \wedge B \wedge C$ with $|\mathbb{T}(A \wedge B \wedge C)| = 1$.

**Case (4)** follows immediately. □

#### 3. Prevalence of different steady states for Boolean tetrahedron networks

**Definition 3.1.** Equilibrium $E$ in tetrahedron network is of type $k+l$ where $k+l = 4, k \geq l$ if $E$ has $k$ entries equal to 1 and $l$ entries equal to 0.

In this section we test the following hypothesis:

**Hypothesis** *Type double positive steady states of the type $(ABcd)$ are more common than single and triple positive steady states $(Abcd), (ABCd)$ in all tetrahedron networks. More specifically, there are more combinations of non-degenerate monotone Boolean functions that exhibit double positive steady states than there are combinations that support single and triple positive steady states $(Abcd), (ABCd)$.*

To simplify the notation we will call any of the six possible doubly positive steady state a $2-2$ steady state and any of the single and triple positive steady states $(Abcd), (ABCd)$ we call a $3-1$ steady state.

##### 3.1. Repressive tetrahedron.

##### 3.2. 3-1 steady states.

Consider without loss of generality equilibrium (0001). We discuss conditions on $f_A, f_B, f_C, f_D$. For all functions $m = (111)$ and $M = (000)$. All functions satisfy non-degeneracy conditions $f_X(m) = 0$ and $f_X(M) = 0$.

- $f_A$**:** The function $f_A$ satisfies condition
$$f_A(001) = 0,$$
which is in group (2) of Theorem 2.1. Therefore there are 2 non-degenerate MBFs.
- $f_B$**:** Function $f_B$ satisfies the same condition.
- $f_C$**:** Function $f_C$ satisfies the same condition.
- $f_D$**:** Function $f_D$ satisfies $f_D(000) = 1$ which falls into group (1) of Theorem 2.1. There are 9 non-degenerate MBFs.

Therefore there are $2 * 2 * 2 * 9 = 72$ non-degenerate collections of MBFs that support 3-1 equilibrium.

##### 3.3. 2-2 steady states.

Consider without loss of generality a steady state (0101).

- $f_A$**:** The function $f_A$ satisfies condition
$$f_A(101) = 0,$$
which is in group (3) of Theorem 2.1. Therefore there are 7 non-degenerate MBFs.

$f_B$: Function $f_B$ satisfies condition

$$f_B(001) = 1,$$

which is in group (3) of Theorem 2.1. Therefore there are 7 non-degenerate MBFs.

$f_C$: Function $f_C$ satisfies condition

$$f_C(011) = 0,$$

which is also in group (3) of Theorem 2.1. Therefore there are 7 non-degenerate MBFs.

$f_D$: Function $f_D$ satisfies condition

$$f_D(010) = 1,$$

which is in group (3) of Theorem 2.1. Therefore there are 7 non-degenerate MBFs.

We conclude that there are $7^4 = 2401$ distinct collections of non-degenerate MBFs supporting 2-2 equilibrium (0101).

The ratio between parameters that support 2-2 equilibria and 3-1 equilibria is

$$\frac{2401}{72} = 33.347.$$

**Hypothesis holds for negative tetrahedron network (1).**

3.4. **Tetrahedron with one positive input into each node.** We assume that the positive edges in the network are

$$A \to B \to C \to D \to A.$$

3.4.1. *3-1 steady states.* Assume without loss of generality 3-1 steady state (0001). We discuss conditions on $f_A, f_B, f_C, f_D$. We note that the function $f_A$ has inputs from BCD in that order, and $D \to A$ is positive. Therefore $m = (110)$ and $M = (001)$. The condition $f_A(001) = 0$ is of the type (4) in Theorem 2.1 and therefore there are non-degenerate MBFs admitting 3-1 equilibrium.

We note that for the symmetric equilibrium $\sigma((0001))$ obtained from (0001) by permutation $\sigma$ there always will be function $f_{\sigma(A)}$ constrained by type (4) constraint.

3.4.2. *2-2 steady states.* Assume without loss of generality 2-2 steady state (0101). We discuss conditions on $f_A, f_B, f_C, f_D$.

$f_A$: Function $f_A$ has inputs from BCD in that order, and $D \to A$ is positive. Therefore $m = (110)$ and $M = (001)$. Since $(101) \in L_M$ condition $f_A(101) = 0$ is in group (2) in Theorem 2.1 and therefore there are 2 non-degenerate MBFs.

$f_B$: Function $f_B$ has inputs from ACD in that order, and $A \to B$ is positive. Therefore $m = (011)$ and $M = (100)$. Since $(001) \in L_m$ condition $f_B(001) = 1$ is in group (2) in Theorem 2.1 and therefore there are 2 non-degenerate MBFs.

$f_C$: Function $f_C$ has inputs from ABD in that order, and $B \to C$ is positive. Therefore $m = (010)$ and $M = (101)$. Since $(011) \in L_m$ condition $f_C(011) = 0$ is in group (3) in Theorem 2.1 and therefore there are 7 non-degenerate MBFs.

$f_D$: Function $f_D$ has inputs from ABC in that order, and $C \to D$ is positive. Therefore $m = (110)$ and $M = (001)$. Since $(010) \in L_M$ condition $f_D(010) = 1$ is in group (3) in Theorem 2.1 and therefore there are 7 non-degenerate MBFs.

Therefore there are 196 distinct collections of nondegenerate MBFs supporting 2-2 equilibrium and no collections suporting 3-1 equilibrium.

**This supports the hypothesis in tetrahedron network with one positive input.**

3.5. **Two positive input tetrahedron.** We assume that the negative edges in the network are

$$A \to B \to C \to D \to A.$$

3.5.1. *3-1 steady states.* Assume without loss of generality 3-1 steady state (0001). For equilibria $\sigma((0001))$ where $\sigma$ is a permutation in $S_4$ the consideration below apply to functions $\sigma(X)$ for $X \in \{A, B, C, D\}$.

We discuss conditions on $f_A, f_B, f_C, f_D$.

$f_A$: Function $f_A$ has inputs from BCD in that order, and $D \to A$ is negative. Therefore $m = (001)$ and $M = (110)$. Since $(101) \in L_m$ condition $f_A(101) = 0$ is in group (3) in Theorem 2.1 and therefore there are 7 non-degenerate MBFs.

$f_B$: Function $f_B$ has inputs from ACD in that order, and $A \to B$ is negative. Therefore $m = (100)$ and $M = (011)$. Since $(001) \in L_M$ condition $f_B(001) = 0$ is in group (2) in Theorem 2.1 and therefore there are 2 non-degenerate MBFs.

$f_C$: Function $f_C$ has inputs from ABD in that order, and $B \to C$ is negative. Therefore $m = (101)$ and $M = (010)$. Since $(011) \in L_T$ condition $f_C(011) = 0$ is in group (2) in Theorem 2.1 and therefore there are 2 non-degenerate MBFs.

$f_D$: Function $f_D$ has inputs from ABC in that order, and $C \to D$ is negative. Therefore $m = (001)$ and $M = (110)$. Since $(010) \in L_B$ condition $f_D(010) = 1$ is in group (2) in Theorem 2.1 and therefore there are 2 non-degenerate MBFs.

This gives 56 collections of non-degenerate MBF supporting 3-1 equilibria.

3.5.2. *2-2 steady states.* Assume an 2-2 steady state (0101). We discuss conditions on $f_A, f_B, f_C, f_D$.

$f_A$: Function $f_A$ has inputs from BCD in that order, and $D \to A$ is negative. Therefore $m = (001)$ and $M = (110)$. Since $(101) \in L_m$ condition $f_A(101) = 0$ is in group (3) in Theorem 2.1 and therefore there are 7 non-degenerate MBFs.

$f_B$: Function $f_B$ has inputs from ACD in that order, and $A \to B$ is negative. Therefore $m = (100)$ and $M = (011)$. Since $(001) \in L_M$ condition $f_B(001) = 1$ is in group (3) in Theorem 2.1 and therefore there are 7 non-degenerate MBFs.

$f_C$: Function $f_C$ has inputs from ABD in that order, and $B \to C$ is negative. Therefore $m = (101)$ and $M = (010)$. Since $(011) \in L_M$ condition $f_C(011) = 0$ is in group (2) in Theorem 2.1 and therefore there are 2 non-degenerate MBFs.

$f_D$: Function $f_D$ has inputs from ABC in that order, and $C \to D$ is negative. Therefore $m = (001)$ and $M = (110)$. Since $(010) \in L_m$ condition $f_D(010) = 1$ is in group (2) in Theorem 2.1 and therefore there are 2 non-degenerate MBFs.

This gives 196 parameters.

**Since** 196 > 56 **this supports the hypothesis for tetrahedron with two positive inputs at each node.**

#### 3.6. Tetrahedron with all positive inputs edges.

3.6.1. *3-1 steady states.* Assume without loss of generality 3-1 steady state (0001). We discuss conditions on $f_A, f_B, f_C, f_D$. For all functions $m = (000)$ and $M = (111)$.

- $f_A$: Function $f_A$ has inputs from BCD in that order. Since $(001) \in L_m$ condition $f_A(101) = 0$ is in group (3) in Theorem 2.1 and therefore there are 7 non-degenerate MBFs.
- $f_B$: Function $f_m$ has inputs from ACD in that order. Since $(001) \in L_m$ condition $f_B(001) = 0$ is in group (3) in Theorem 2.1 and therefore there are 7 non-degenerate MBFs.
- $f_C$: Function $f_C$ has inputs from ABD in that order. Since $(001) \in L_m$ condition $f_C(001) = 0$ is in group (2) in Theorem 2.1 and therefore there are 7 non-degenerate MBFs.
- $f_D$: Function $f_D$ has inputs from ABC in that order. Since $(000) = m$ condition $f_D(000) = 1$ is in group (4) in Theorem 2.1 and therefore there are no non-degenerate MBFs.

There are no collections of non-degenerate MBFs that support with 3-1 steady states.

3.6.2. *2-2 steady states.* Assume without loss of generality 2-2 steady state (0101). We discuss conditions on $f_A, f_B, f_C, f_D$. For all functions $m = (000)$ and $M = (111)$.

- $f_A$: Function $f_A$ has inputs from BCD in that order. Since $(101) \in L_M$ condition $f_A(101) = 0$ is in group (2) in Theorem 2.1 and therefore there are 2 non-degenerate MBFs.
- $f_B$: Function $f_B$ has inputs from ACD in that order. Since $(001) \in L_m$ condition $f_B(001) = 1$ is in group (2) in Theorem 2.1 and therefore there are 2 non-degenerate MBFs.
- $f_C$: Function $f_C$ has inputs from ABD in that order. Since $(011) \in L_M$ condition $f_C(011) = 0$ is in group (2) in Theorem 2.1 and therefore there are 2 non-degenerate MBFs.
- $f_D$: Function $f_D$ has inputs from ABC in that order. Since $(010) \in L_m$ condition $f_D(010) = 1$ is in group (2)in Theorem 2.1 and therefore there are 2 non-degenerate MBFs.

Therefore there are 16 collections that support 2-2 steady states.

**This supports the hypothesis for positive tetrahedron network.**

### 4. Assessing relative strengths of mutual connections

In this section we consider the following question for repressive tetrahedron. For either an 3-1 steady state or a 2-2 steady state, given a pair of nodes $A, B$ is there a pattern in relative strength of connections $A \to B$ vs. $B \to A$?

Recall that for TTr network we have $m = (111)$ and $M = (000)$. Therefore we assess the strength of repressive connection $A \to B$ by the number of value 1 in the corresponding monotone Boolean function $f$ i.e. the size of the truth set $|\mathbb{T}(f)|$.

For pair of nodes $A, B$ mutually connected by repressing edges we will count number of pairs of monotone Boolean functions $(f_A, f_B)$ that support the equilibrium for which $|\mathbb{T}(f_A)| > |\mathbb{T}(f_B)|$, the number where$|\mathbb{T}(f_A)| = |\mathbb{T}(f_B)|$, and the number of pairs where $|\mathbb{T}(f_A)| < |\mathbb{T}(f_B)|$.

Due to tetrahedron $S_4$ symmetry there for each equilibrium the only one number that must be computed is the number between the node that is ON, and node that is OFF at the equilibrium. The identity of the nodes within each category does not matter.

4.1. **3-1 steady state.** Without loss of generality we consider steady state (0001) that was also considered in section 3.2. Without loss consider nodes $A$ and $D$. The steady state is supported by two MBFs $f_A$ at node $A$ one with $|\mathbb{T}(f)| = 1$ and one with $|\mathbb{T}(f)| = 3$. On the other hand, the steady state is supported at the node $D$ (which is ON) by all 9 MBFs $f_D$ with the size of the truth set $1, 3, 3, 3, 4, 5, 5, 5, 7$, respectively. Taking all combinations between these groups we find that

- in 1 out of 18 cases we have $|\mathbb{T}(f_A)| > |\mathbb{T}(f_D)|$;
- in 4 out of 18 cases we have $|\mathbb{T}(f_A)| = |\mathbb{T}(f_D)|$;
- in 13 out of 49 cases we have $|\mathbb{T}(f_A)| < |\mathbb{T}(f_D)|$.

Therefore among the parameters that support the equilibrium (0001) (of the type $(abcD)$); the frequency of parameters where $|\mathbb{T}(f_A)| < |\mathbb{T}(f_D)|$ is larger than frequency of parameters where $|\mathbb{T}(f_A)| > |\mathbb{T}(f_D)|$. This supports the conclusion that among these the strength of repression from $A$ to $D$ is generally smaller that repression from $D$ to $A$.

4.2. **2-2 steady state.** Without loss of generality we consider steady state (0101) that was considered in section 3.3 and consider nodes $A$ and $D$ which are OFF and ON respectively. The steady state is supported by 7 non-degenerate MBFs $f_A$ at node $A$ which have at least two zeroes and hence $|\mathbb{T}(f)_A| \leq 6$, and by 7 non-degenerate MBFs $f_D$ at the node $D$ with $|\mathbb{T}(f_D)| \geq 2$. This results for $f_A$ with $|\mathbb{T}(f_A)| \in \{1, 3, 3, 3, 4, 5, 5\}$, since one of the functions with $|\mathbb{T}(f_A)| = 5$ is not compatible with the condition $f_A(101) = 0$. Similarly, the 7 functions $|\mathbb{T} f_D| \in \{3, 3, 4, 5, 5, 5, 7\}$. For the 49 combinations of these functions we find that

- in 8 out of 49 cases we have $|\mathbb{T}(f_A)| > |\mathbb{T}(f_D)|$.
- in 13 out of 49 cases we have $|\mathbb{T}(f_A)| = |\mathbb{T}(f_D)|$;
- in 28 out of 49 cases we have $|\mathbb{T}(f_A)| < |\mathbb{T}(f_D)|$;

Therefore among the parameters that support the equilibrium (0101) (of the type $(aBcD)$); the frequency of parameters where $|\mathbb{T}(f_A)| < |\mathbb{T}(f_D)|$ is larger than frequency of parameters where $|\mathbb{T}(f_A)| > |\mathbb{T}(f_D)|$. This again supports the conclusion that among these parameters the strength of repression from $A$ to $D$ is generally smaller that repression from $D$ to $A$.

These results support conclusions of Fig. 4 of the main text.
