## Supplementary Figures S1-S8 for "Multistability and predominant double-positive states in a four node mutually repressive network: a case study of Th1/Th2/Th17/T-reg differentiation"

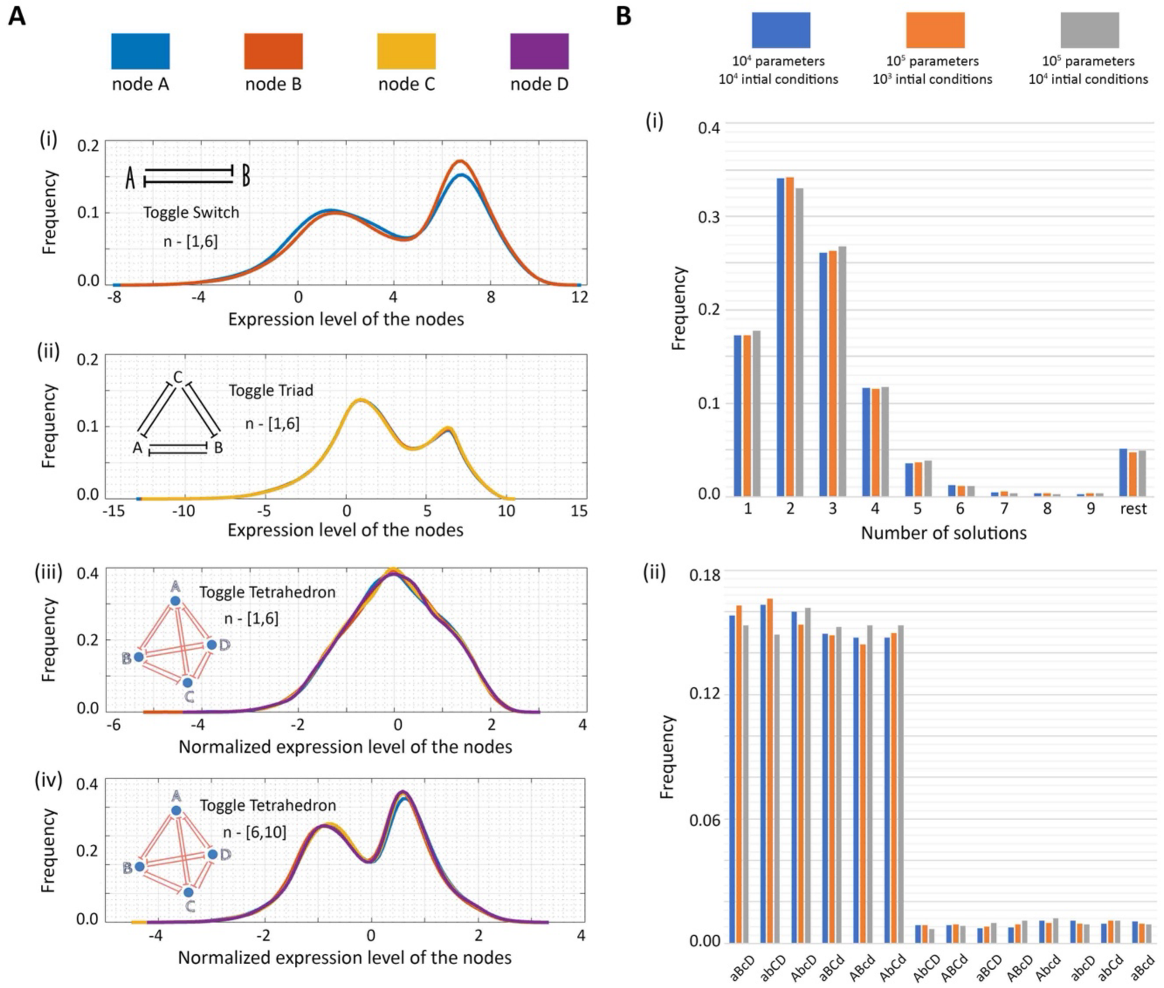


*Figure S1: (A) (i) Node expression values obtained from RACIPE results of Toggle Switch network motif plotted as a distribution to display the modality (here, bimodal) (ii) Same as (i) but for Toggle Triad (iii) Same as (i) but for Toggle Tetrahedron with low values of the hill coefficient (iv) Same as (iii) but for high values of the hill coefficient. (B) (i) Frequency of monostable, bistable, tristable, tetrastable, pentastable solutions and the rest in the toggle tetrahedron for RACIPE simulations of different number of initial conditions and parameter sets generated. (ii) Frequency of all states taken together for the same cases scenarios as in (i).*

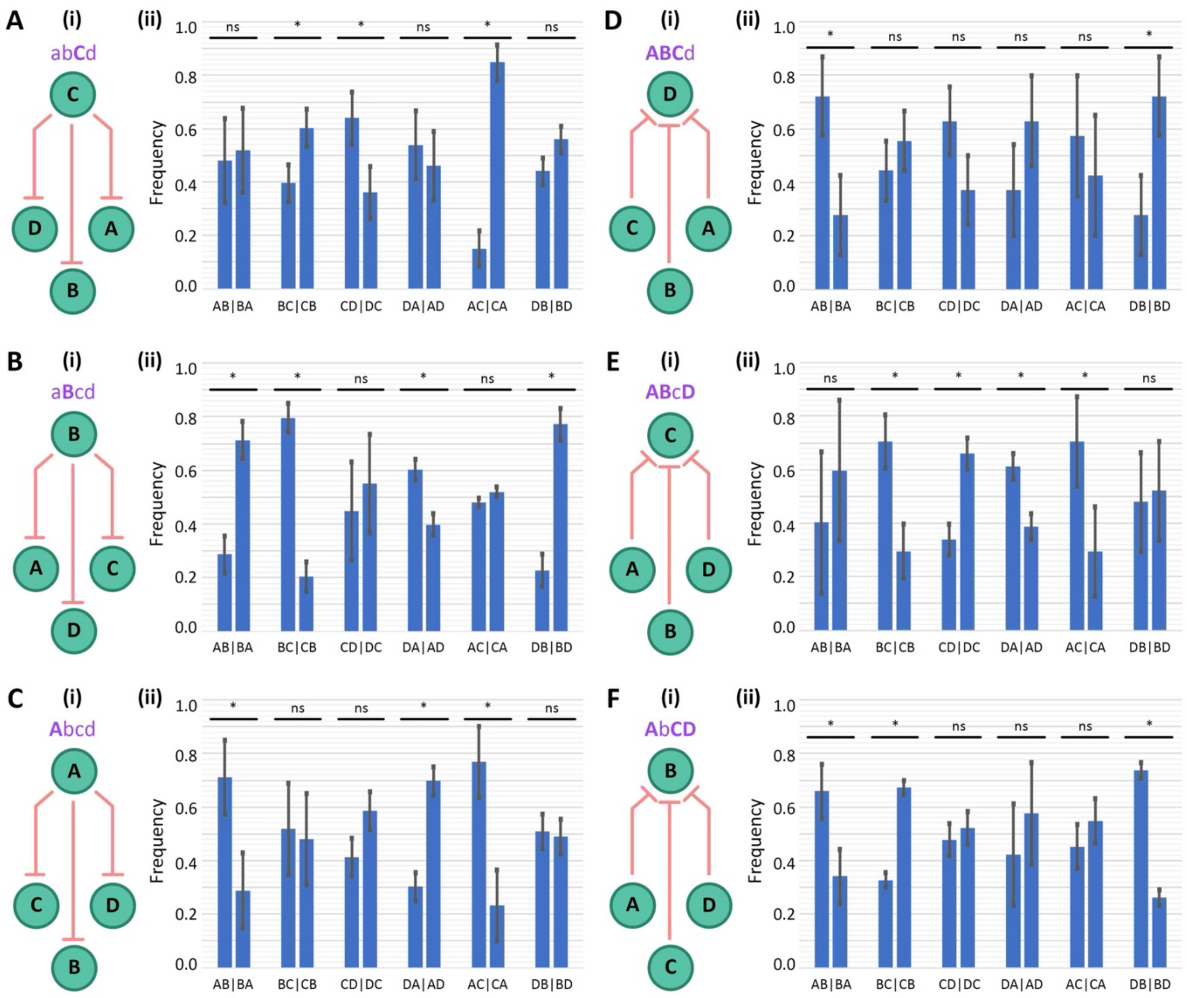


*Figure S2: RACIPE - Link Strength Analysis of monostable states for toggle tetrahedron. (A) (i) Schematic showing the links expected to be stronger than their counterparts for the sate {abCd} (ii) Frequency of dominance of all twelve coupled links for the case of monostable {abCd}. (B) Same as (A) but for the monostable state – {aBcd}. (C) Same as (A) but for the monostable state – {Abcd}. (D) Same as (A) but for the monostable state – {ABCd}. (E) Same as (A) but for the monostable state – {ABcD}. (F) Same as (A) but for the monostable state – {AbCD}.*

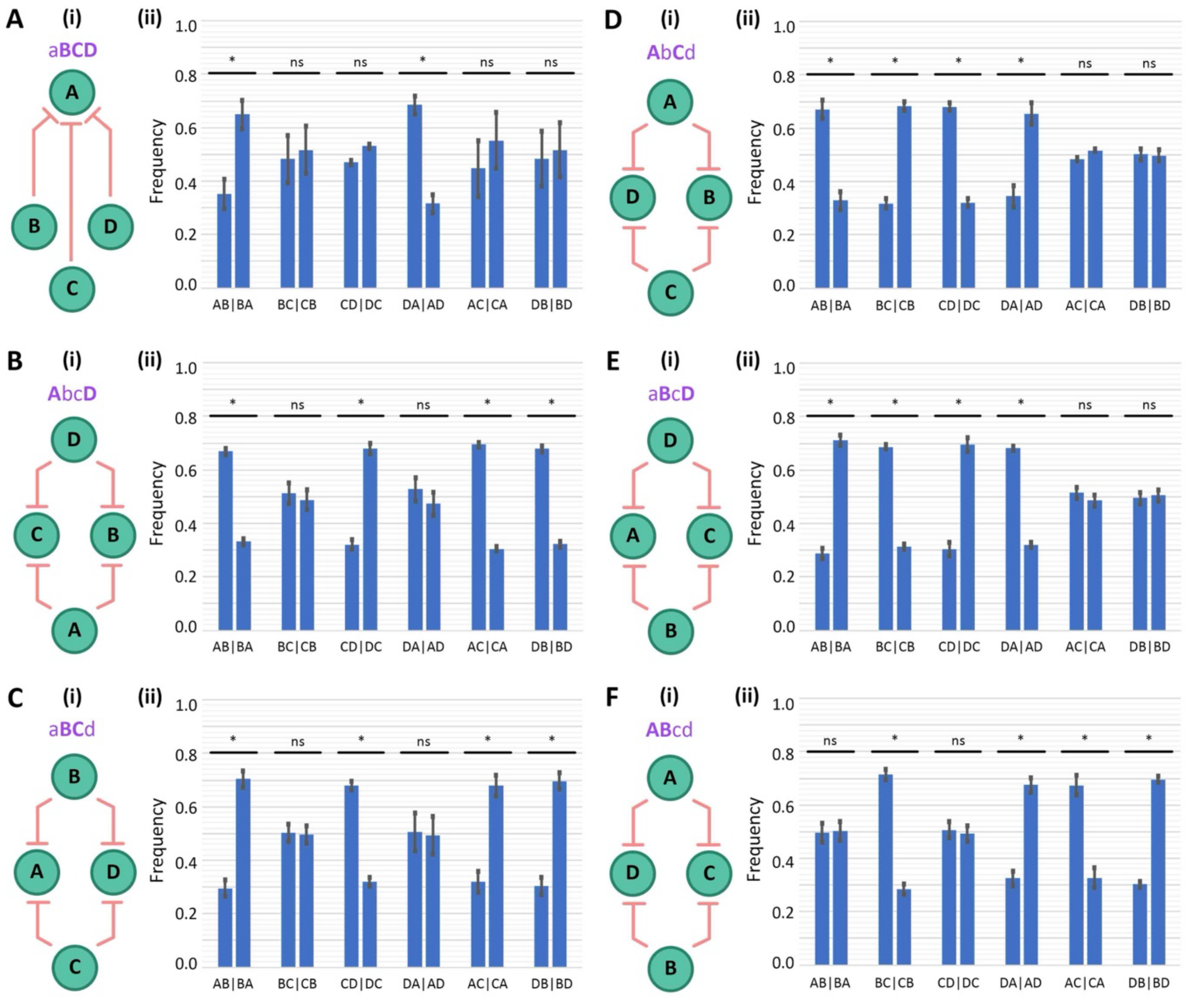


*Figure S3:* *RACIPE - Link Strength Analysis of monostable states for toggle tetrahedron. (A) (i) Schematic showing the links expected to be stronger than their counterparts for the sate {aBCD} (ii) Frequency of dominance of all twelve coupled links for the case of monostable {aBCD}. (B) Same as (A) but for the monostable state – {AbcD}. (C) Same as (A) but for the monostable state – {aBCd}. (D) Same as (A) but for the monostable state – {AbCd}. (E) Same as (A) but for the monostable state – {aBcD}. (F) Same as (A) but for the monostable state – {ABcd}.*

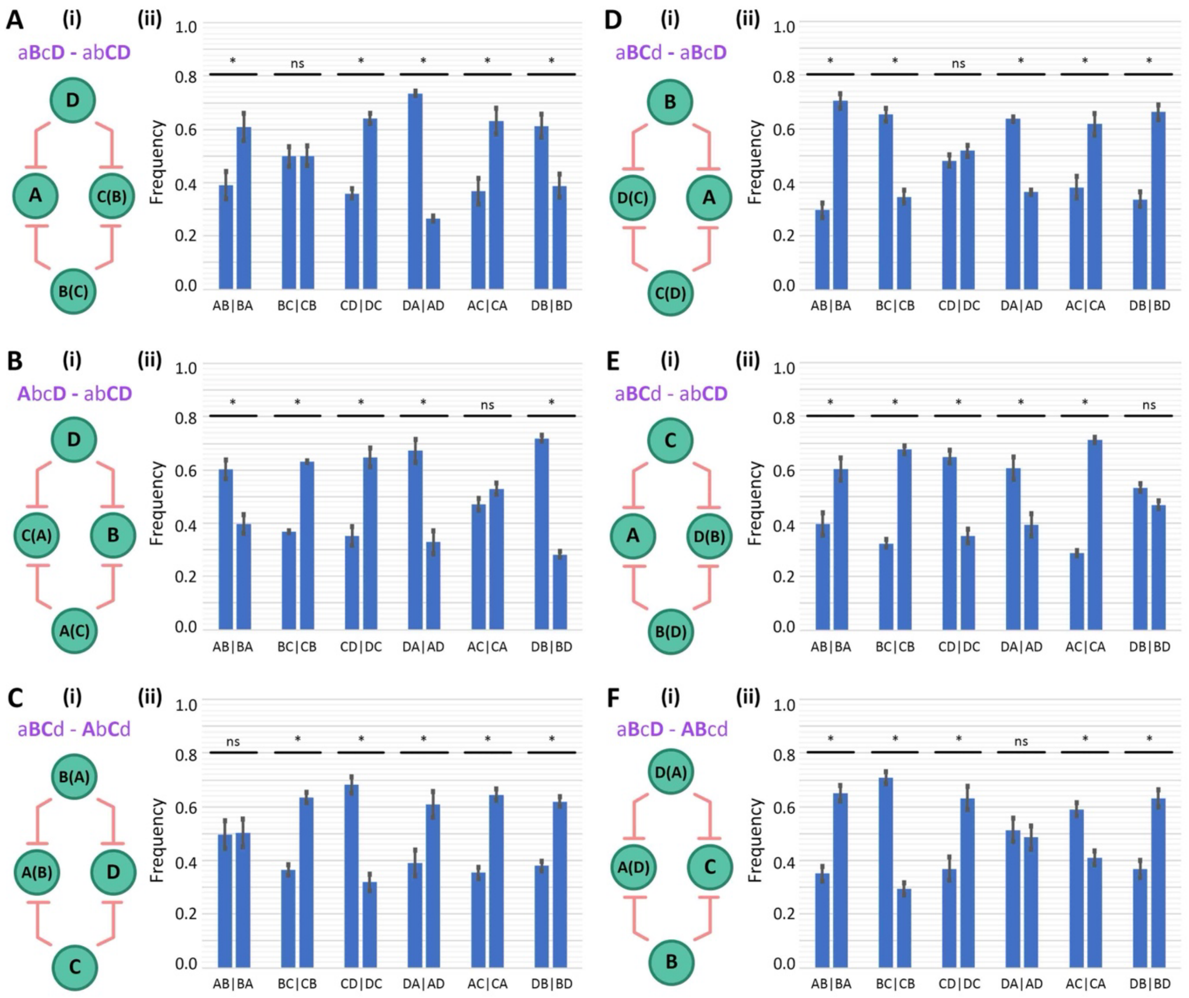


*Figure S4:* RACIPE- Link Strength Analysis of bistable states for toggle tetrahedron. (A) (i) Schematic showing the links expected to be stronger than their counterparts for the non-mirror bistable sate {aBcD, abCD} (ii) Frequency of dominance of all twelve coupled links for the case of non-mirror bistable sate {aBcD, abCD}. (B) Same as (A) but for – {AbcD, abCD}. (C) Same as (A) but for – {aBCd, AbCd}. (D) Same as (A) but for – {aBCd, aBcD}. (E) Same as (A) but for – {aBCd, abCD}. (F) Same as (A) but for – {aBcD, ABcd}.

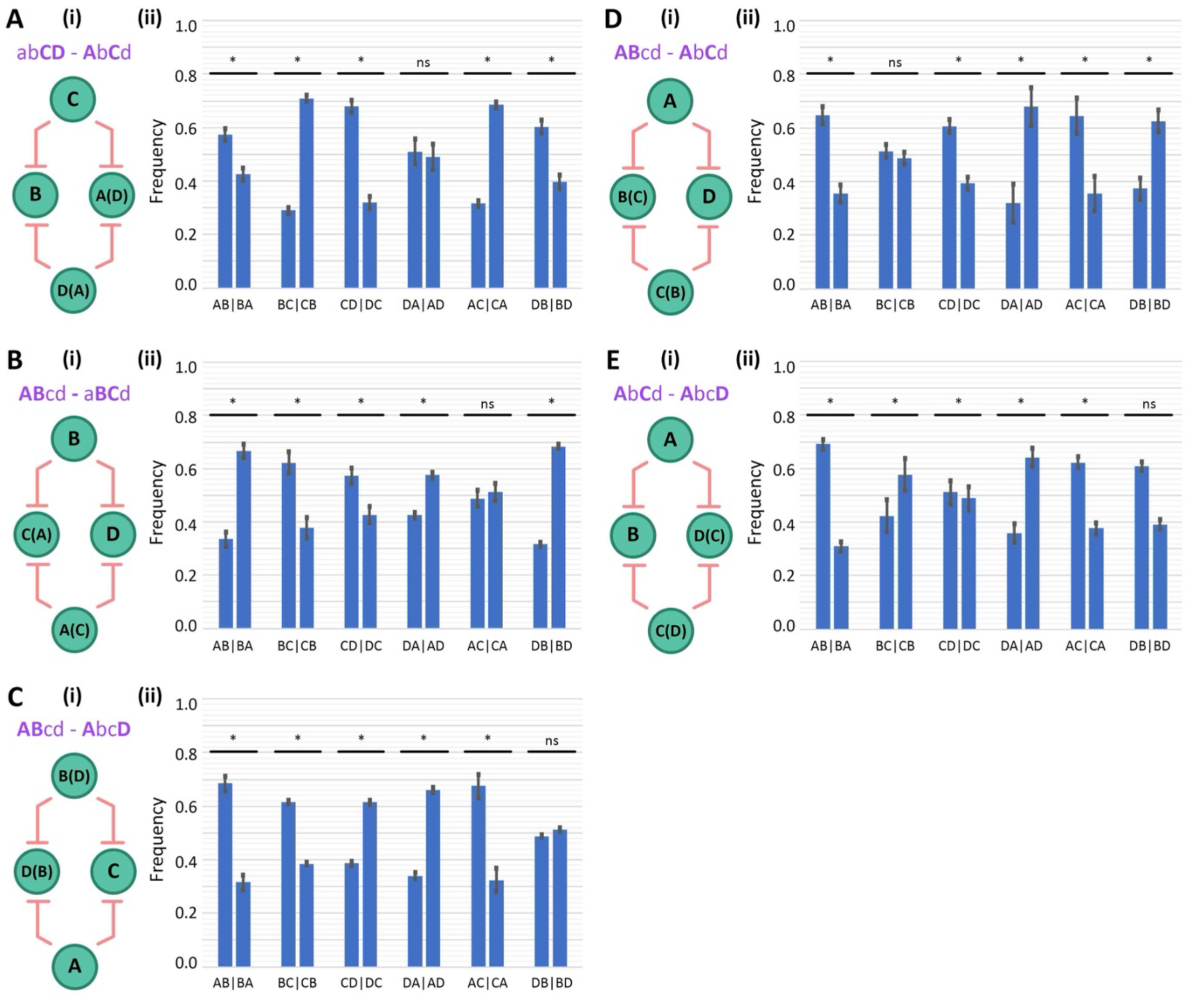


*Figure S5:* RACIPE- Link Strength Analysis of bistable states for toggle tetrahedron. (A) (i) Schematic showing the links expected to be stronger than their counterparts for the non-mirror bistable sate {abCD, AbCd} (ii) Frequency of dominance of all twelve coupled links for the case of non-mirror bistable sate {abCD, AbCd}. (B) Same as (A) but for – {ABcd, aBCd}. (C) Same as (A) but for – {ABcd, AbcD}. (D) Same as (A) but for – {ABcd, AbCd}. (E) Same as (A) but for – {AbCd, AbcD}.

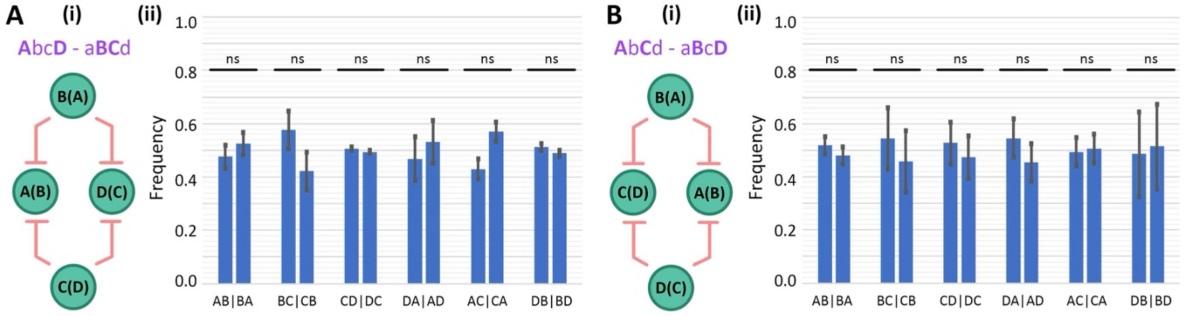


*Figure S6:* RACIPE- Link Strength Analysis of bistable states for toggle tetrahedron. (A) (i) Schematic showing the links expected to be stronger than their counterparts for the mirror bistable sate {AbcD, aBCd} (ii) Frequency of dominance of all twelve coupled links for the case of mirror bistable sate {AbcD, aBCd}. (B) Same as (A) but for – {AbCd, aBcD}.

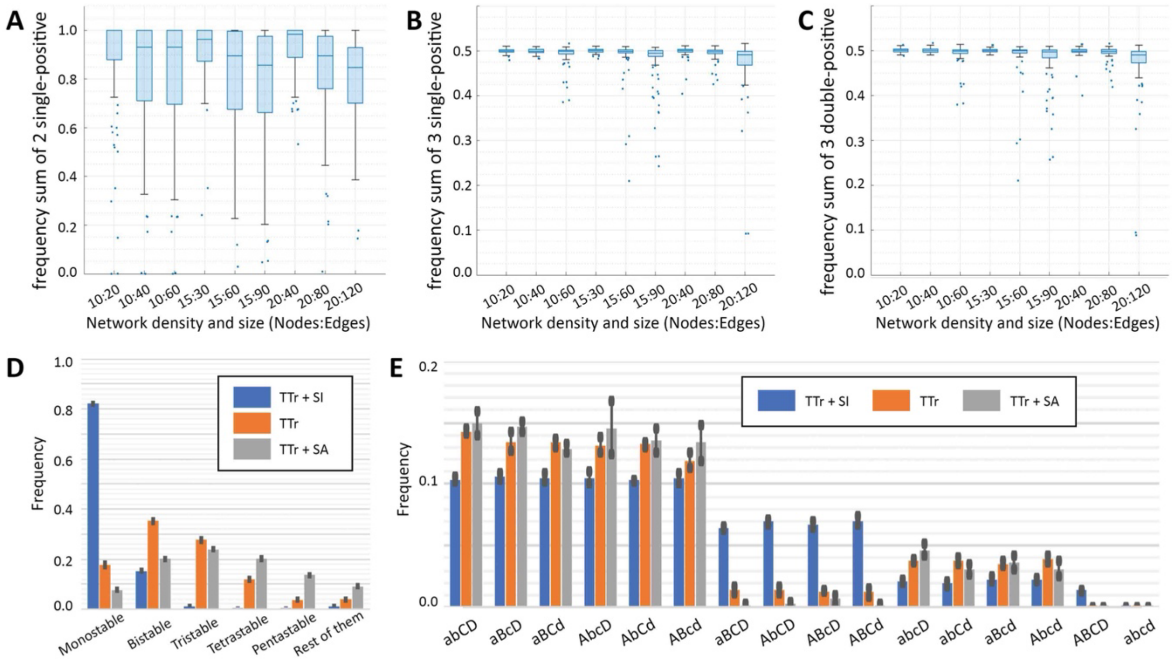


*Figure S7:* (A) Sum of frequencies of the two 'single-positive' states – {Ab, aB} for Boolean simulations of toggle switch embedded in large networks of varying sizes and densities. (B) Sum of frequencies of the three 'single-positive' states – {Abc, aBc, abC} for Boolean simulations of toggle triad embedded in large networks of varying sizes and densities. (C) Same as (B) but for the three 'double-positive' states – {ABc, aBC, AbC}. (D) State frequency distribution for TTr, TTr+SI and TTr+SA. (E) Frequency distribution of states when only the monostable states are considered for TTr, TTr+SI and TTr+SA.

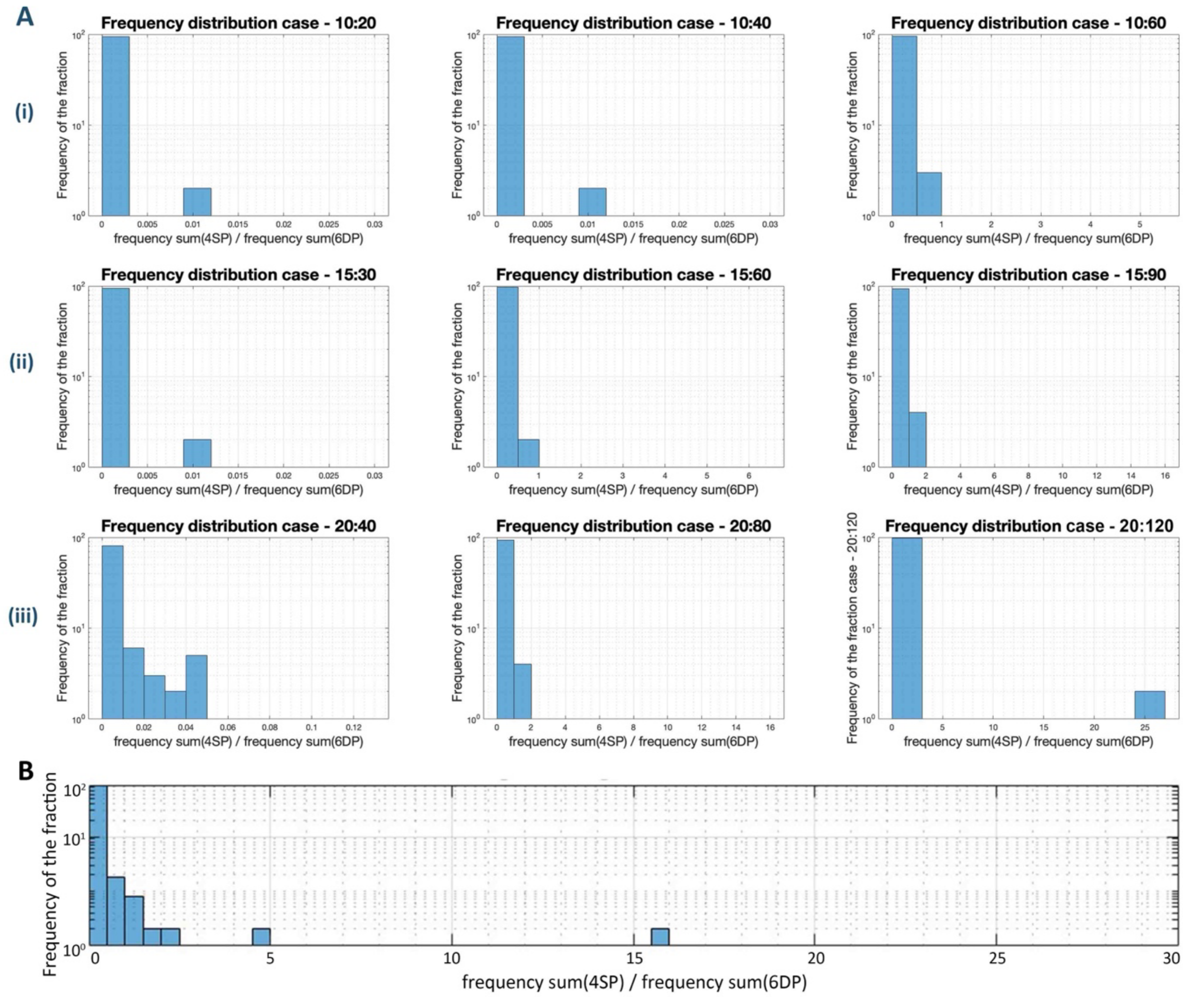


*Figure S8:* (A) (i) Histograms of fraction of sum frequency of the 4 single-positive (SP) states over the sum frequency of the 6 double-positive (DP) states calculated for all 100 random networks of size – 10 nodes and densities of 2, 4 and 6 times the number of nodes (as mentioned in the subplot titles). (ii) Same as (i), but for network size – 15 nodes. (iii) Same as (i), but for network size – 20 nodes. (B) Same as (i), but considering all network sizes and densities together.
